# The alternative polyadenylation regulator CFIm25 promotes macrophage differentiation and activates the NF-κβ pathway

**DOI:** 10.1101/2024.09.03.611136

**Authors:** Srimoyee Mukherjee, Atish Barua, Luyang Wang, Bin Tian, Claire L. Moore

**Affiliations:** Department of Developmental, Molecular, and Chemical Biology, Tufts University School of Medicine, Boston, Massachusetts 02111, USA; The Wistar Institute, Philadelphia, Pennsylvania 19104, USA

**Keywords:** alternative polyadenylation, monocyte, macrophage, CFIm25, cell cycle, NUDT21

## Abstract

Macrophages are required for our body’s development and tissue repair and protect against microbial attacks. We previously reported a crucial role for regulation of mRNA 3’-end cleavage and polyadenylation (C/P) in monocyte to macrophage differentiation. The CFIm25 subunit of the C/P complex showed a striking increase upon differentiation of monocytes with Phorbol Myristate Acetate, suggesting that it promotes this process. To test this hypothesis, CFIm25 was overexpressed in two different monocytic cell lines, followed by differentiation. Both cell lines showed a significant increase in macrophage characteristics and an earlier slowing of the cell cycle. In contrast, depletion of CFIm25 hindered differentiation. Cell cycle slowing upon CFIm25 overexpression was consistent with a greater decrease in the proliferation markers PCNA and cyclin D1, coupled with increased 3’UTR lengthening of cyclin D1 mRNA. Since choice of other poly(A) sites could be affected by manipulating CFIm25, we identified additional genes with altered use of poly(A) sites during differentiation and examined how this changed upon CFIm25 overexpression. The mRNAs of positive regulators of NF-κB signaling, TAB2 and TBL1XR1, and NFKB1, which encodes the NF-κB p50 precursor, underwent 3’UTR shortening that was associated with increased protein expression compared to the control. Cells overexpressing CFIm25 also showed elevated levels of phosphorylated NF-κB-p65 and the NF-κB targets p21, Bcl-XL, ICAM1 and TNF-α at an earlier time and greater resistance to NF-κB chemical inhibition. In conclusion, our study supports a model in which CFIm25 accelerates the monocyte to macrophage transition by promoting alternative polyadenylation events which lead to activation of the NF-κB pathway.

## 1 Introduction

Monocytes and macrophages are members of the mononuclear phagocyte system, a component of innate immunity (1). Monocytes proliferate in response to infection and injury, and after rushing to the site of interest, mature into active macrophages that can combat invading pathogens or promote tissue repair (2). In order to manipulate macrophages to be more effective at these roles, we need to better understand how gene expression is regulated during their differentiation.

Alternative polyadenylation (APA) is a widespread mechanism that regulates mRNA and protein isoform expression by determining the position where poly(A) is added to the 3’ end of mRNA transcripts (3–5). APA often results in mRNA isoforms with the same coding sequence but different 3’ UTRs lengths, with downstream effects on mRNA stability, translatability, localization, and RNA-seeded protein/protein interactions, while intronic APA alters the coding sequence. We have recently documented alternative polyadenylation (APA) as an important phenomenon in the complex process of monocyte-macrophage differentiation and showed a role for the Cleavage/Polyadenylation (C/P) protein CstF64 in promoting this differentiation (6). However, many global APA regulators have been identified (7, 8), and it is likely that other proteins besides CstF64 will influence macrophage differentiation through APA. A possible candidate is the mRNA 3’-processing factor CFIm, which is known to be a principal regulator of 3′ UTR length in many cell types (9–15). In our previous study, we have found that compared to other C/P proteins, the CFIm25 (CPSF5/NUDT21) subunit showed an increase after induction of macrophage differentiation at a time when the first signs of differentiation were starting to appear (6). We hypothesized that it might be a significant player in the process of monocyte to macrophage differentiation.

CFIm25 binds specifically to a UGUA motif as a dimer (16, 17) and associates with either the CFIm68 (CPSF6) or CFIm59 (CPSF7) subunit to form an active CFIm complex (18). Mechanistically, CFIm25 exerts its effect on cell fate by inducing a widespread switch of APA patterns in hundreds of transcripts (14). Multiple studies have revealed that CFIm25 levels affect the proliferation of many types of tumor cells, with CFIm25 being mostly tumor-suppressive, in part by causing lengthening of transcripts of the cell cycle gene CCND1 (14, 19, 20). CFIm25 also has antifibrotic properties, and its down regulation amplifies lung, skin, and liver fibrosis (21–23). CFIm25 blocks the generation of pluripotent stem cells (15) and is needed for the differentiation of embryonic stem cells into the ectodermal lineage. Its depletion hinders differentiation of neutrophils from a myeloid progenitor cell line (15), but the affected molecular pathways and altered APA events in this context were not identified.

In the current study, we report that overexpression of CFIm25 accelerates macrophage differentiation, as assessed by changes in the rate of cell attachment, cell cycle progression, and expression of the macrophage-specific CD38. Importantly, CFIm25 regulates expression of genes in the NF-κB pathway, which plays a major role in maturation of monocytes to macrophages and protecting these cells from apoptosis (24–27). The APA and protein level of two positive regulators of the NF-κB pathway, TAB2 and TBL1XR1 (TBLR1), were modulated by CFIm25. TAB2 is an activator of the TAK1 kinase, which is required for the IL-1 induced activation of NF-kβ (28), while TBL1XR1 is a NF-κB transcriptional cofactor that promotes transcription of NF-κB target genes (29–31). We also found poly(A) site (PAS) switching in mRNAs of NFKB1, encoding the precursor of the p50 DNA-binding subunit of the NF-kβ complex. These findings establish CFIm25 as a positive regulator of the monocyte to macrophage transition that acts in part by activating the NF-κB pathway.

## 2 Results

### 2.1 Macrophage differentiation is accompanied by enhanced CFIm25 expression

To study the impact of alternative polyadenylation (APA) in monocyte to macrophage differentiation, we used two well-established monocytic cell lines, HL-60 and THP-1 (32, 33), which can be differentiated to macrophages by treatment with PMA (34–36). Treatment of both cell lines was done for a time course (0-24 h), and as observed previously (6, 37), these cells, which normally grow in suspension, displayed increased adherence to the culture dish and no longer proliferated after PMA treatment (Fig. 1A and B). By 24 h, most cells showed other macrophage-like phenotypes such as increased cell size and appearance of filopodia (data not shown).

**Fig 1.**
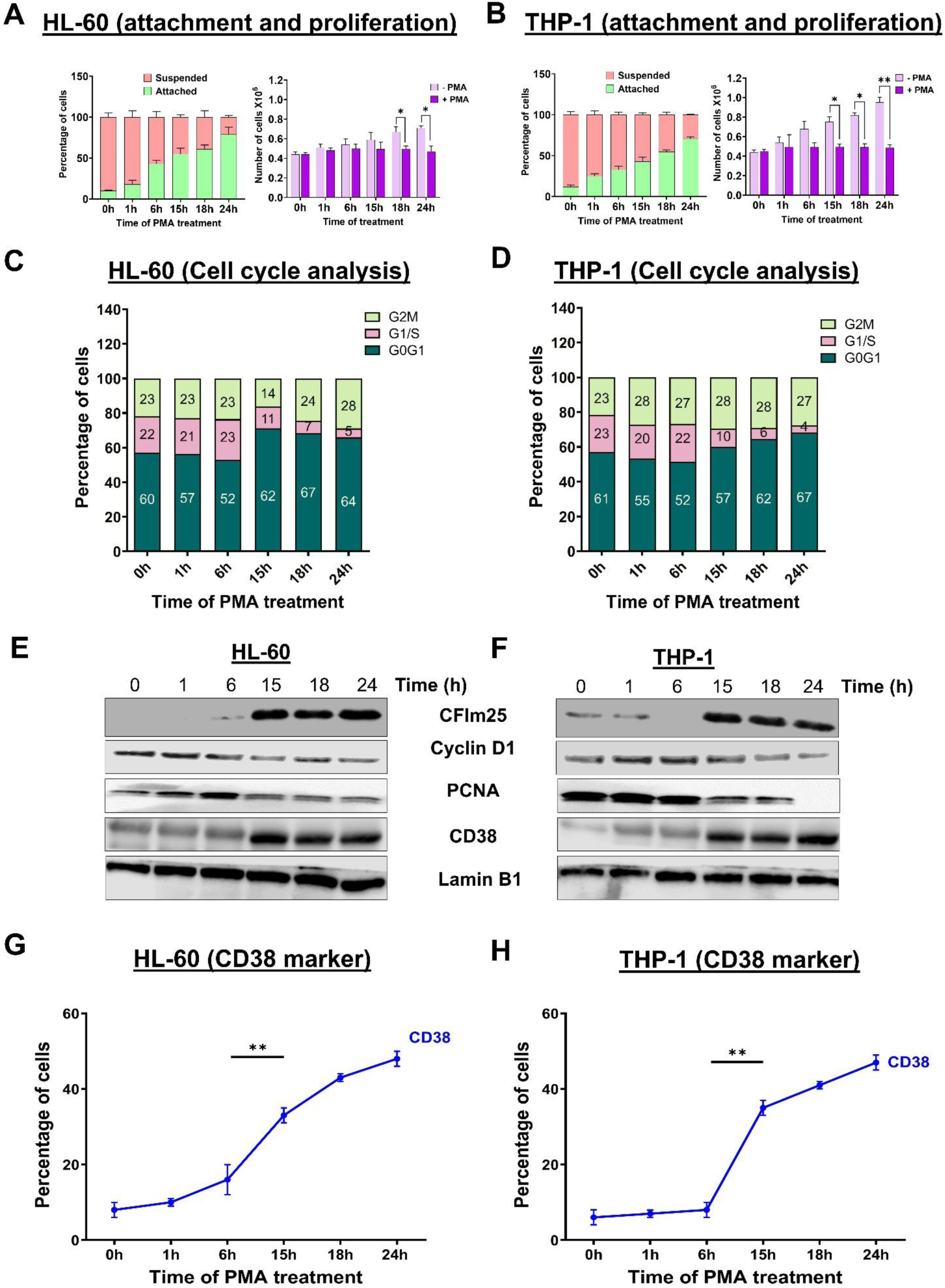
PMA treatment leads to differentiation and enhanced CFIm25 expression. **(A & B) Attachment and proliferation assays in (A) HL-60 and (B) THP-1 cells.** For the attachment assay (left), cells were treated with 3 nM PMA for 0, 1, 6, 15, 18 and 24 h and live cells were visualized and counted by the Trypan blue exclusion assay. The graph presents the percentage of cells that are suspended or attached. For the proliferation assay (right), the number of cells before and after differentiation depicts the absolute cell number in millions, compared to normal cell proliferation in the absence of PMA. The figure represents mean ± SE from three independent experiments. P value <0.05 was considered significant, where * = P ≤ 0.05; **= P ≤ 0.01. **(C & D) Cell cycle analysis for the HL-60 (C) and THP-1 (D) time courses.** Cell cycle progression was determined by PI staining followed by flow cytometry at the indicated times after PMA treatment. Quantitative measurement of cell cycle phase is presented as a stacked bar graph. Numbers in each column represent the percentage of cells in the specified phase. Data is representative of at least three biological replicates **(E & F) Western blots for markers in HL-60 (E) and THP-1 (F) cells.** Western blot analysis of the macrophage marker CD38, the cell cycle marker cyclin D1, the proliferation marker PCNA and the C/P protein CFIm25 in cells treated with PMA for the indicated times, where GAPDH serves as the loading control. **(G & H) CD38 levels by flow cytometry of (G) HL-60 and (H) THP-1 cells.** The expression of CD38 over time was analyzed by flow cytometry and is shown as % CD38+ cells. Data was obtained from at least three biological replicates. The figure represents mean ± SE from three independent experiments. P value <0.05 was considered significant, where **** = P ≤ 0.0001

Cells usually slow down their cell cycle progression while differentiating into distinct cell types (38). By examining the cell cycle distribution of HL-60 and THP-1 cells during differentiation with flow cytometry, we found that for both cell types, the response to PMA involved slowing of cell cycle progression from 15 h onwards, with fewer cells in the G1/S phase (Fig. 1C and D, and Supplementary Fig. 1A and B). Consistent with decreased proliferation, the levels of Proliferating Cell Nuclear Antigen (PCNA), a well-known proliferation marker, and Cyclin D1 (CCND1), a protein which modulates the transition from G1 to S phase both decreased in expression over the differentiation time course of HL-60 and THP-1 cells (Fig. 1E-F). Macrophage differentiation is also associated with increased expression of macrophage markers (39). By western blot analysis, we observed an increase in the macrophage marker CD38 (40) (Fig. 1E-F). Flow cytometry analysis showed that CD38 surface expression was also increased (Fig. 1G-H and Supplementary Fig. 1C-D).

We recently showed that the cellular changes occurring during macrophage differentiation were accompanied by an overall increase in the expression of the C/P complex proteins, including CFIm25 (6). In agreement with our earlier work, PMA treatment in the present study increased the levels of CFIm25 in both HL-60 and THP-1 cells (Fig. 1E-F). As described in the Introduction, the CFIm25 protein is known to promote alternative polyadenylation in many cellular transitions, and our goal in this study was to determine the role of CFIm25 in macrophage differentiation.

### 2.2 CFIm25 overexpression promotes macrophage differentiation and expedites the cell cycle exit

To find out if overexpression of CFIm25 in PMA-treated cells would accelerate the appearance of differentiation phenotypes, we constitutively overexpressed CFIm25 in both HL-60 and THP-1 cells with the help of lentiviral constructs (Fig. 2A and B). There was a marked increase in attachment at the 6h time with no change in cell viability in CFIm25 overexpressing cells compared to controls (Supplementary Fig. 2A and B). An earlier and stronger decrease in Cyclin D1 and PCNA further indicated enhanced differentiation (Fig. 2A and B). In accordance with this finding, cell cycle analysis for the cells overexpressing CFIm25 showed slowing of cell cycle progression earlier in the differentiation time course compared to the controls for both cell lines (Fig. 2C and D and Supplementary Fig. 2C and D). An enhanced differentiation is also evident from an earlier increase (at 6h) in the macrophage marker CD38 as demonstrated by western blot (Fig. 2A and B) and flow cytometry analysis (Fig. 2E-F and Supplementary Fig. 2E-F). In summary, overexpression of CFIm25 slows cell cycle progression as evident from flow cytometry and repression of PCNA and Cyclin D1 and enhances expression of a surface marker that distinguishes monocytes from macrophages. These findings indicate that the normal increase in CFIm25 during differentiation expedites the transition of monocytes to macrophages.

**Fig 2.**
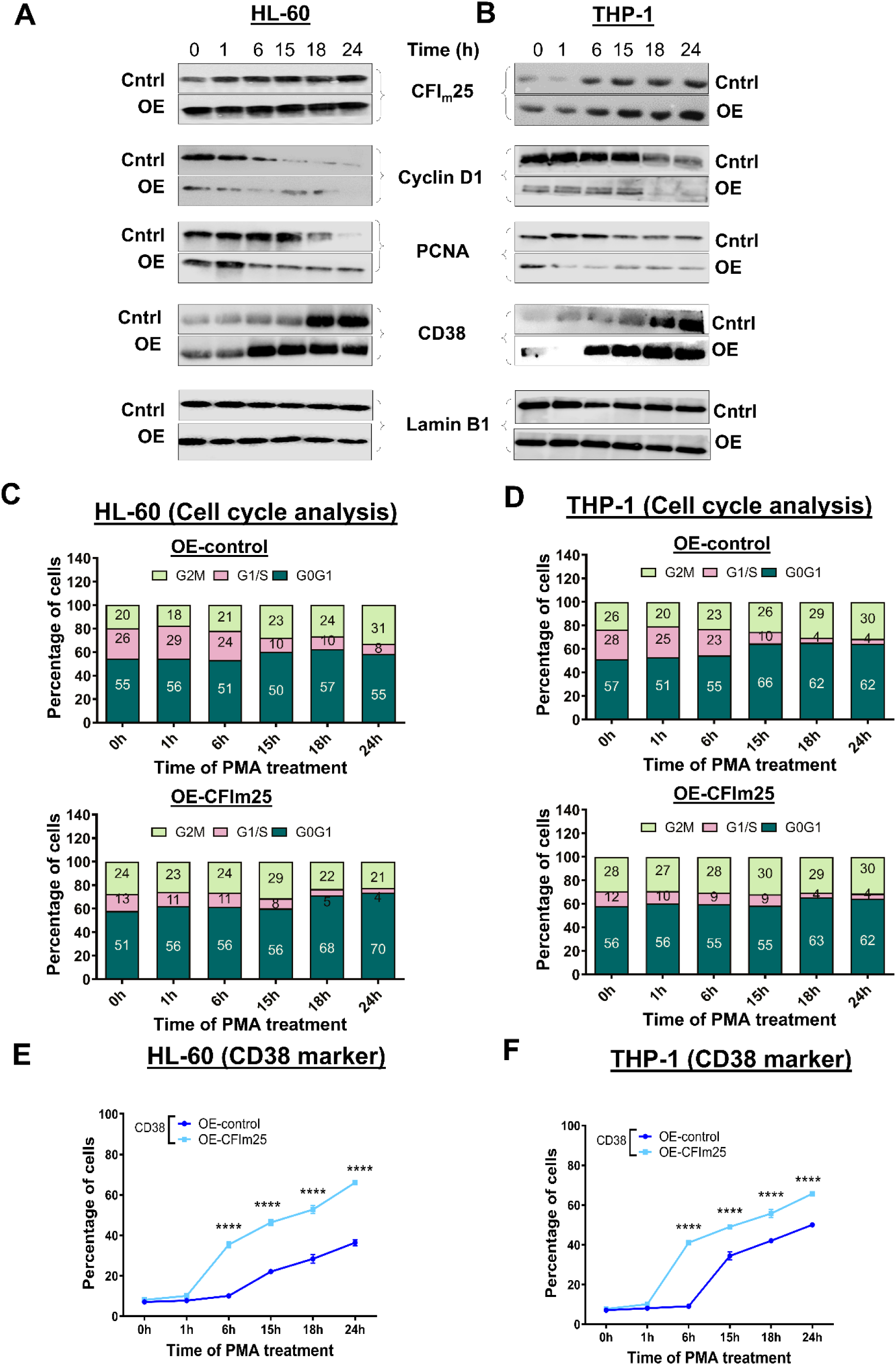
CFIm25 overexpression promotes the differentiation to macrophages and expedites the cell cycle exit. **(A & B) Western blots in HL-60 (A) and THP-1 (B) cells.** Western blot analysis of CFIm25, cyclin D1, PCNA and CD38 for cells overexpressing CFIm25 (OE) with respect to control (Cntrl), at the indicated times of PMA treatment. GAPDH serves as the loading control. **(C & D) Cell cycle analysis for (C) HL-60 or (D) THP-1 cells overexpressing CFIm25 and treated with PMA.** Cell cycle analysis was performed as described in Fig. 1. **(E & F) CD38 levels by flow cytometry of (E) HL-60 and (F) THP-1 control cells or cells overexpressing CFIm25.** The expression of CD38 over time was analyzed is shown as % CD38+ cells. Data is representative of at least three biological replicates and is plotted as mean ± SE. P value <0.05 was considered significant, where * = P ≤ 0.05; **= P ≤ 0.01; *** = P ≤ 0.001.

### 2.3 CFIm25 knockdown delays differentiation to macrophages

To determine if CFIm25 depletion has effects on differentiation to macrophages opposite to that of overexpression, expression of CFIm25 in HL-60 and THP-1 cells was knocked down by transfecting pooled siRNAs against this C/P protein. There was a considerable decrease in CFIm25 protein expression, indicating efficient knockdown (Fig. 3A and B, upper panel). After PMA treatment and CFIm25 knockdown, these cells displayed decreased attachment but no change in viability (Supplementary Fig. 3A and B). For both cell lines, cell cycle analysis after CFIm25 knockdown and PMA treatment showed no drop in the number of cells in G1/S phase for HL-60 cells and a slower drop for THP-1 cells (Fig. 3C and D, and Supplementary Fig. 3C and D), and no decline in the levels of Cyclin D1 and PCNA (Fig. 3A and B). In addition, by western blot, the macrophage marker CD38 fails to increase over the time course in comparison to the control cells (Fig. 3A and B). By flow cytometry analysis, expression of the macrophage marker CD38 on the cell surface also did not increase with CFIm25 KD at the initial (6h) time (Fig. 3E-F and Supplementary Fig. 3E-F). These findings indicate that depleting CFIm25 leads to a strong delay in differentiation of monocytes to macrophages.

**Fig 3.**
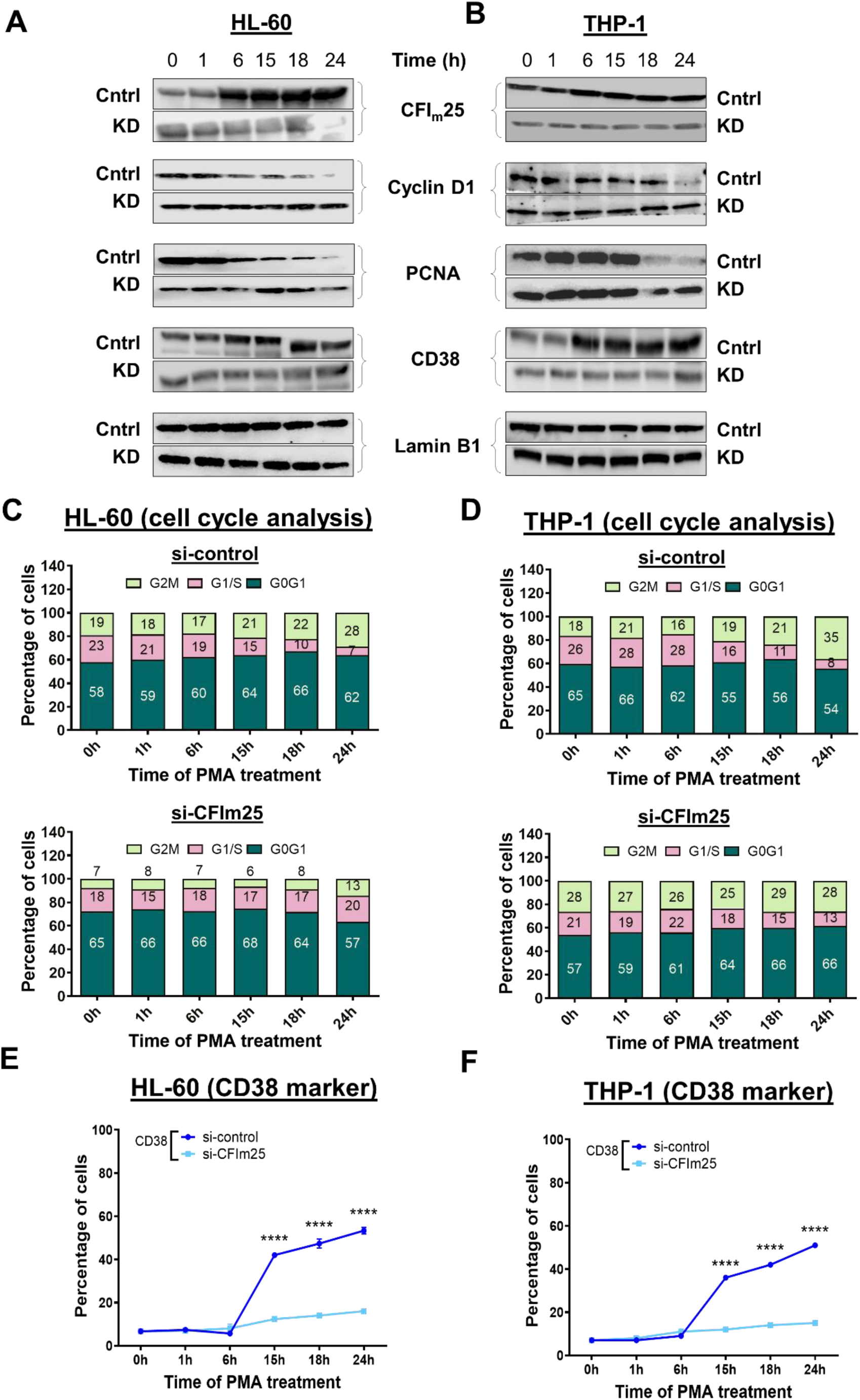
CFIm25 knockdown delays slowing of cell cycle and differentiation to macrophages. **(A & B) Western blots for markers in HL-60 (A) and THP-1 (B) cells.** Western blot analysis of CFIm25, cyclin D1, PCNA and CD38 at the indicated times of PMA treatment for cells knocked down with siRNA against CFIm25 (KD) with respect to si-control (Cntrl). GAPDH serves as the loading control for both the cells. **(C & D) Cell cycle analysis after knockdown of CFIm25 in (C) HL-60 and (D) THP-1 cells treated with PMA.** Cell cycle analysis was performed as described in Fig. 1. **(E & F) CD38 levels by flow cytometry of (E) HL-60 and (F) THP-1 control cells depleted of CFIm25.** The expression of CD38 over time is shown as % CD38+ cells. Data is representative of at least three biological replicates and is plotted as mean ± SE. P value <0.05 was considered significant, where * = P ≤ 0.05; **= P ≤ 0.01; *** = P ≤ 0.001.

### 2.4 Alternative polyadenylation of CCND1 may contribute to the CFIm25-mediated cell cycle regulation

CCND1 (Cyclin D) mRNA is subject to APA and usage of its proximal PAS has been reported to accelerate the cell cycle and promote cell proliferation of HEK293T cells (19). In another study, distal PAS usage of CCND1 was reported to decrease after CFIm25 knockdown in lung cancer cells (20). Since differentiated monocytes and those overexpressing CFIm25 displayed decreased protein levels of cyclin D, we wanted to investigate whether the CCND1 APA status could contribute to this decrease. We first determined the total level of CCND1 mRNA expression in HL-60 and THP-1 cell lines using primers designed to amplify the exon 2–3 junction (Supplementary Table 2). CCND1 mRNA decreases in the control cells between 0 and 18 hours and decreases even further upon CFIm25 overexpression (Fig. 4A and 4B). Next, we measured APA changes by a PCR-based method, where long isoforms were detected with a primer pair just upstream of the mapped distal PAS (Supplementary Table 2), and the ratio of long/total mRNA expression was used to determine the change in usage of the distal site (Fig. 4C and 4D). Using this strategy, we found significantly increased usage of the distal sites, i.e., formation of longer isoforms upon CFIm25 overexpression during macrophage differentiation. In contrast, CFIm25 knockdown resulted in an increased expression coupled with significant shortening of CCND1 transcript during the course of differentiation (Fig. 4E-H, D). Several studies have found that the longer isoform of CCND1 is associated with decreased cyclin D1 protein in HEK293T and cancer cell lines (19, 20, 41). Thus, the lengthening that we observe in CCND1 mRNAs could explain the decrease in cyclin D1 protein and contribute to faster cell cycle exit during macrophage differentiation.

**Fig 4.**
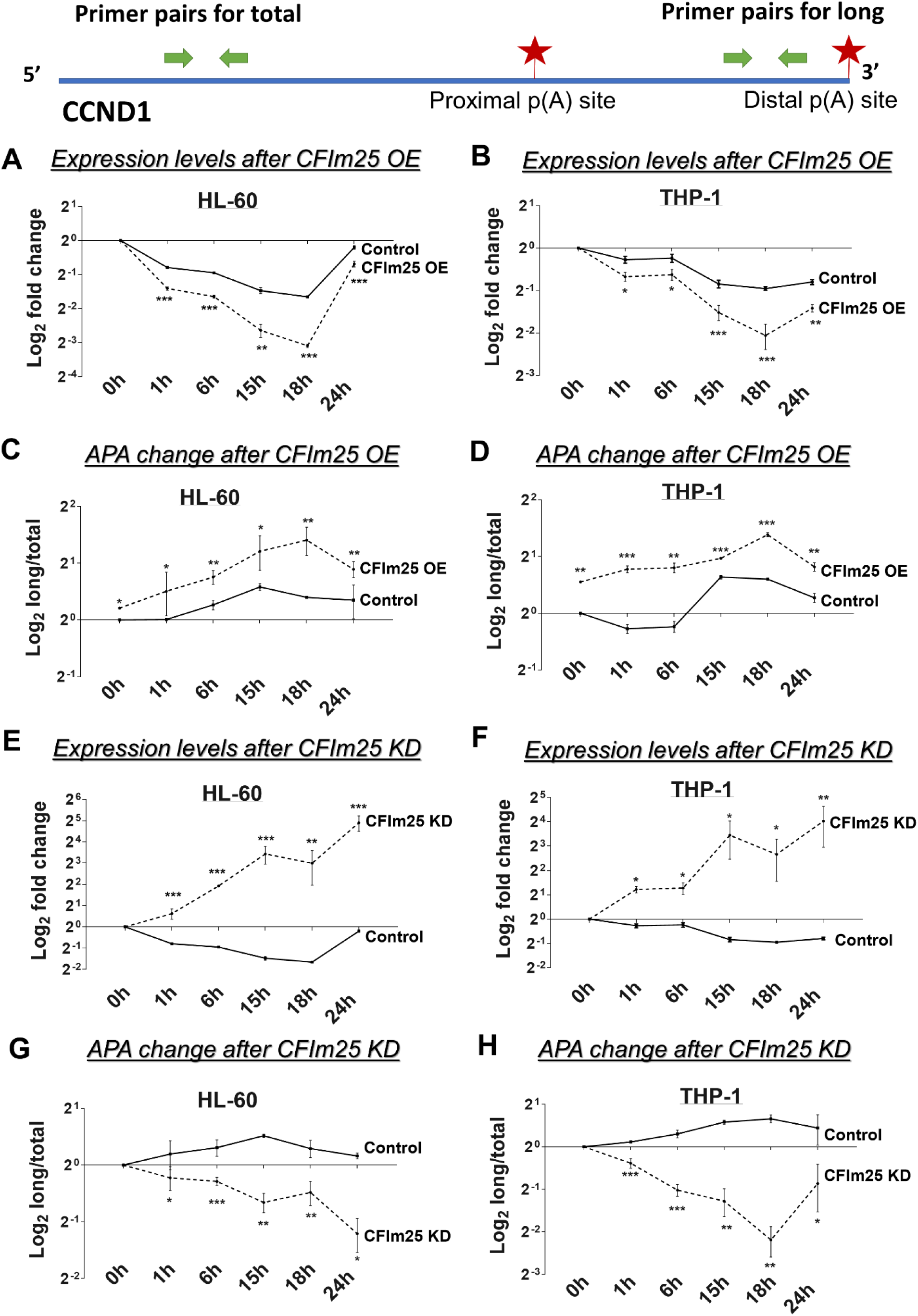
APA and expression of CCND1 after CFIm25 manipulation. Real-time quantitative PCR (RT-qPCR)–based analysis of the expression and distal PAS usage of the cell cycle gene CCND1. The RT-qPCR analysis used two pairs of primers, where one represents the total transcript level and the other primer pair targets sequences just upstream of the distal PAS to detect long transcripts. Positions of primer pairs for each gene are provided in the upper panel and sequences in Supplementary Table 2. **(A & B)** Fold changes (log_2_) in expression of this gene during the 0-24 h time course post PMA treatment of HL-60 and THP-1 cells which are either transfected with control or constructs overexpressing CFIm25, where values are normalized to ACTB mRNA. **(C and D)** Changes in ratio of long to total transcript levels of control and CFIm25 overexpressing cells. **(E and F)** Expression changes after knocking down CFIm25 compared to controls. **(G and H)** Changes in ratio of long to total transcript levels of control and CFIm25-depleted cells. Both bar graphs represent mean ± SE from three independent experiments. P value <0.05 was considered significant, where * = P ≤ 0.05; **= P ≤ 0.01; *** = P ≤ 0.001.

### 2.5 Global effect of CFIm25 overexpression on alternative polyadenylation during macrophage differentiation

To determine the effects of CFIm25 on other PAS usage during macrophage differentiation, we used the QuantSeq 3’ mRNA-sequencing method (42), which can provide information on differential APA events. PolyA+-selected RNA samples from HL-60 cells overexpressing CFIm25 and control cells after 6 h of PMA treatment were sequenced and analyzed to identify changes in isoform usage. We selected the 6 h timepoint because at this time, the control cells are entering a transition phase marked by slowing of the cell cycle and changes in the expression of the surface marker CD38 (Fig. 2C and 2E). We classified significant APA events as those where P ≤ 0.05 and found two genes to be lengthened (NFYC and UCK2) and eight genes to be shortened (CHMP4B, FERMT3, IREB2, OAZ1, SRI, TAB2, TBL1X1 and TM9SF2). Detailed outputs for these APA events are supplied in Supplementary Table 3. Genome browser plots showing the positions and changes in PAS usage for these genes are shown in Supplementary Fig. 4.

We used a PCR-based method (10, 23) to confirm a change in mRNA isoforms from the genes that differed in their 3’ ends. Using a coding sequence primer pair (Supplementary Table 2), we found that while total mRNAs of the lengthened NFYC and UCK2 genes were decreased, all others (CHMP4B, FERMT3, IREB2, OAZ1, SRI, TAB2, TBL1XR1 and TM9SF2) were increased upon CFIm25 overexpression (Fig. 5A, upper panel). Transcripts extending beyond the proximal PASs were detected with a primer pair just upstream of mapped distal PASs (Supplementary Table 2), and the ratio of long isoform/total transcripts was used to determine change in usage relative to the control cells (Fig. 5A, lower panel). Using this strategy, we found an increased use of distal p(A) sites of NFYC and UCK2 mRNAs (lengthening) and a decreased usage (shortening) for the other genes after CFIm25 overexpression (Figure 5A, bottom panel). These results are in accordance with the QuantSeq data (Supplementary Figure 4). Furthermore, we observed similar directional changes in protein levels as seen with the mRNA expression (Figure 5B), in agreement with studies suggesting that shortened 3′ UTRs often lead to an increased stability of the mRNAs and an increased protein output (3–5). We found similar changes in mRNA expression, PAS usage, and protein levels for these genes upon CFIm25 overexpression in THP-1 cells, except that SRI transcripts were not shortened (Supplementary Fig. 5A and B). The possible roles of the genes undergoing CFIm25-induced APA are described later in the Discussion.

**Fig 5.**
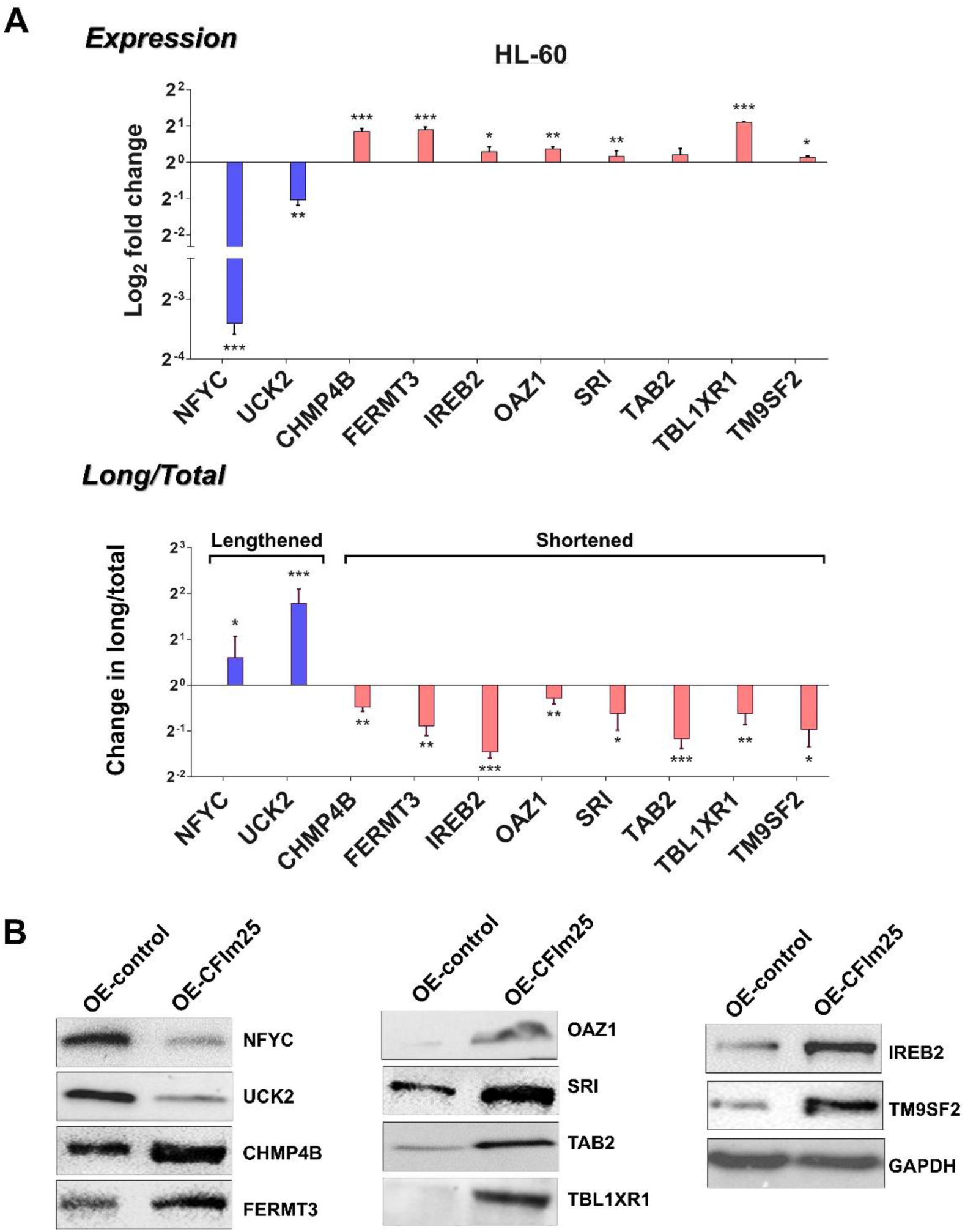
CFIm25 overexpression results in APA changes of specific genes and altered protein expression in PMA-treated HL-60 cells. **(A) APA and expression analysis of CFIm25 target genes.** Real-time quantitative PCR (RT-qPCR)–based analysis of the changes in expression and distal PAS usage of genes in HL-60 cells identified by the QuantSeq method as undergoing transcript lengthening (NFYC, UCK2) or shortening (CHMP4B, FERMT3, IREB2, OAZ1, SRI, TAB2, TBL1XR1 and TM9SF2) upon CFIm25 overexpression. Analysis was performed as described in Fig. 4. P value <0.05 was considered significant, where * = P ≤ 0.05; **= P ≤ 0.01; *** = P ≤ 0.001. **(B) Protein levels of genes with APA changes.** Western blot analysis of proteins encoded by the shortened and lengthened genes in HL-60 cells overexpressing CFIm25 compared to control. GAPDH serves as the loading control.

### 2.6 CFIm25 affects the differentiation and cell cycle changes through the NF-κB pathway

NF-κB has been implicated as a master regulator of monocytes and macrophages (24), and activation of this pathway could be beneficial for acute myeloid leukemia (AML) patients by inducing terminal myeloid differentiation (43). Of the targets we examined that undergo alterations in their 3’ UTR lengths, TAB2 and TBL1XR1 directly contribute to the NF-κB pathway (28–31). NFKB1, encoding the p50 subunit of NF-κB, is also reported to exhibit APA site switching during the antiviral immune response (44). While the two cell lines showed differences in the timing or magnitude of changes in mRNA and protein during the differentiation time course response to CFIm25 overexpression, both HL-60 and THP-1 cells showed consistent shortening of the 3’ UTRs of NFKB1, TAB2 and TBL1XR1 mRNAs (Fig. 6A and 6C) that was coupled with increased mRNA (Fig. 6B and 6D) and more and/or earlier protein expression (Fig. 7).

**Fig 6.**
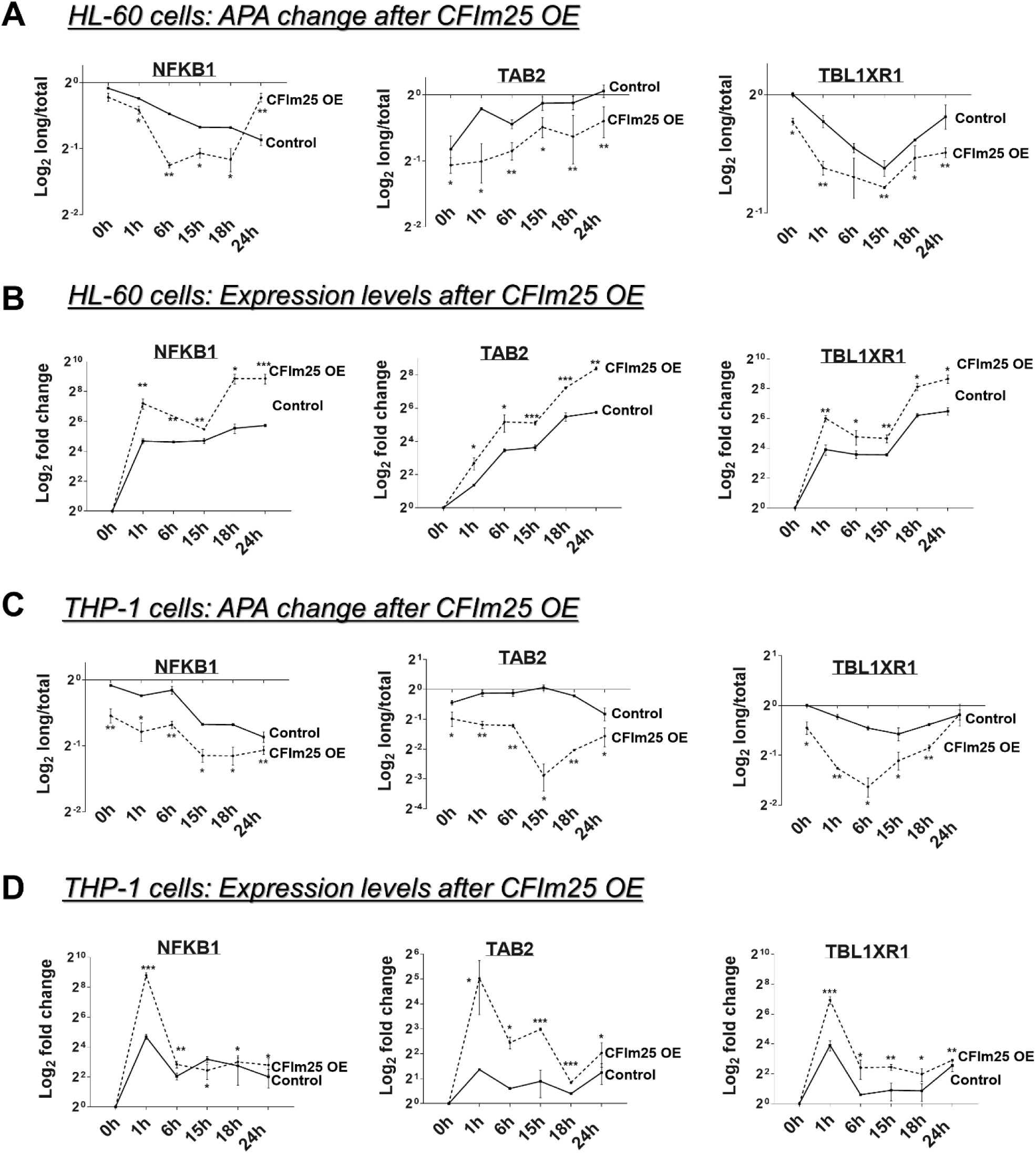
CFIm25 affects mRNA expression and APA of effectors of the NF-κB pathway. Real-time quantitative PCR (RT-qPCR)–based analysis of distal PAS usage and the expression of NFKB1, TAB2 and TBL1XR1 after overexpression of CFIm25 and treatment with PMA for 0-24h in HL-60 (A, B) and THP-1 (C, D) cells. The analysis was performed as described in Fig. 4.

**Fig 7.**
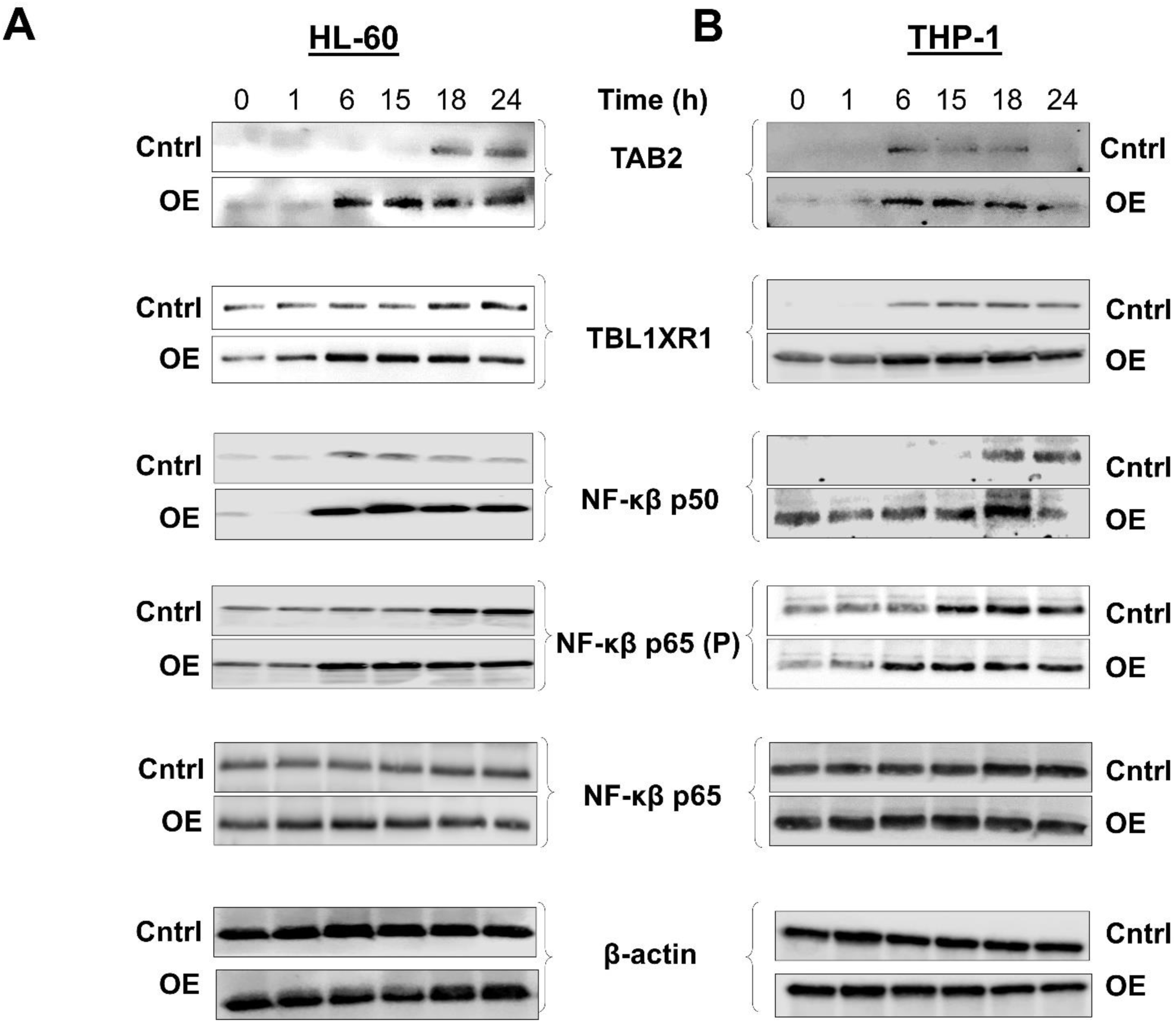
CFIm25 overexpression alters levels of the effectors of the NF-κB pathway. Western blot analysis of the indicated proteins during the differentiation time course for HL-60 (A) and THP-1 (B) cells upon overexpression of CFIm25 and time course treatment with PMA (0-24 h). GAPDH serves as a loading control.

We next examined the expression of the NF-κB p65 (RelA) subunit and downstream targets by western blot. An earlier increase in the levels of phosphorylated p65 after CFIm25 overexpression indicates that the NF-κB pathway is hyperactivated (Fig. 7). Important downstream targets of the NF-κB pathway (45), such as p21, the adhesion molecule ICAM1, and the anti-apoptotic factor Bcl-XL, were increased in protein levels (Fig. 8A-B). As it is well known that NF-κB mediates the induction of cytokines in monocytes and macrophages (24), we also measured the levels of TNF-α by ELISA. CFIm25 overexpression caused a strong increase in TNF-α (Fig. 8C and 8D). Similar results were found in both HL-60 and THP-1 cells.

**Fig 8.**
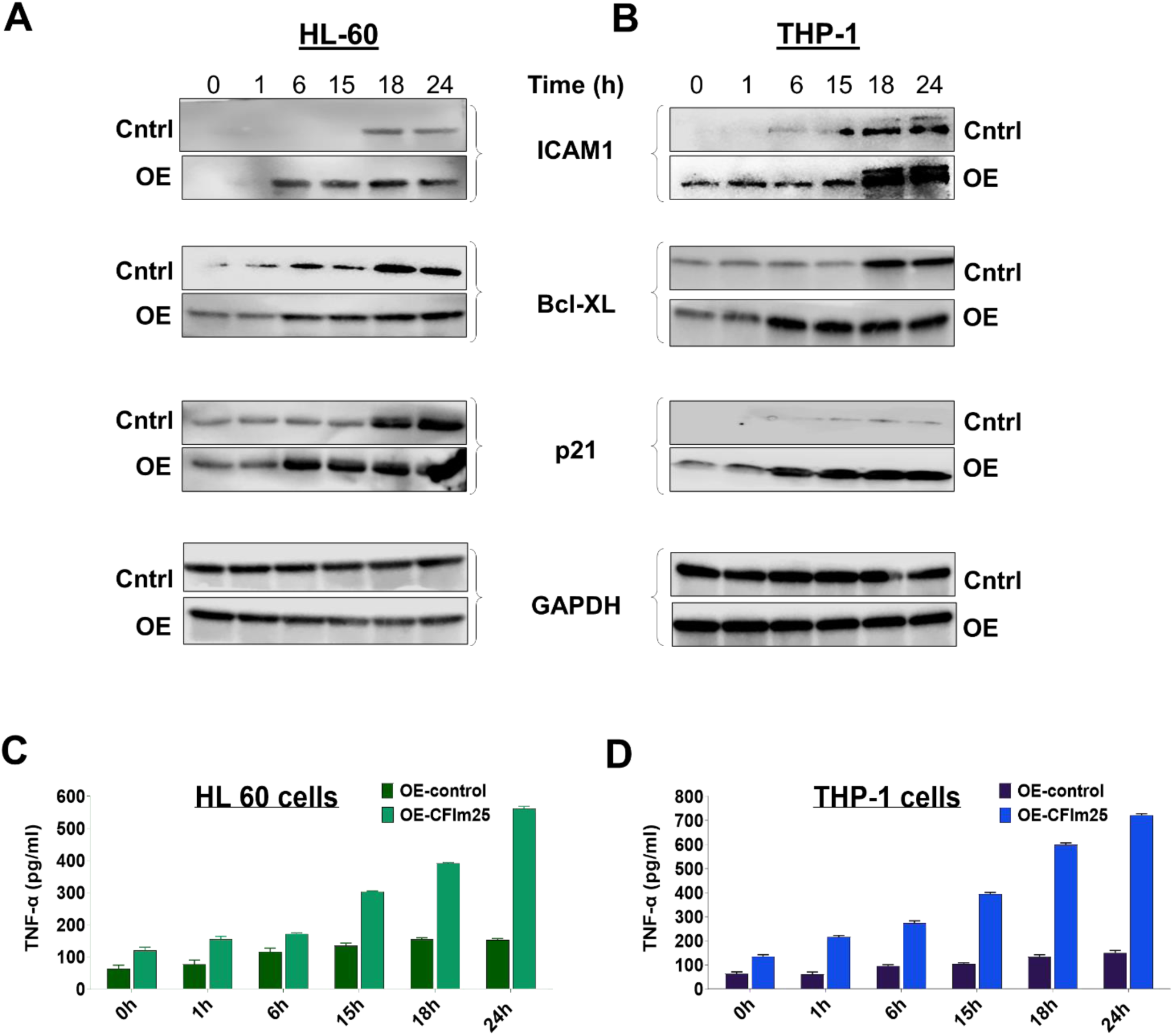
CFIm25 overexpression affects the targets of the NF-κB pathway. Western blot analysis of the indicated proteins during the differentiation time course for HL-60 (A) and THP-1 (B) cells upon overexpression of CFIm25 and treatment with PMA (0-24 h). GAPDH serves as a loading control. **(C & D) ELISA for TNF-α.** ELISA of TNF-α in HL-60 (C) and THP-1 (D) cell supernatants upon overexpression of CFIm25 with respect to controls after induction of differentiation.

To further assess NF-κB activation, we treated control or CFIm25 overexpressing HL-60 or THP-1 cells with the NF-κB inhibitor BAY11-7082 (43) and subjected them to attachment, viability and flow cytometric analysis for macrophage markers. As expected, treatment with BAY 11-7082 greatly reduced the attachment of the PMA-treated cells without affecting their viability (Fig. 9 and 9B) and decreased the percentage of cells expressing the macrophage markers CD38 and CD11b (Fig. 9C and 9D), indicating impeded transition of monocytes to macrophages. In contrast, cells overexpressing CFIm25 showed a higher level of attachment and marker expression after BAY 11-7082 treatment compared to the control cells treated with the inhibitor, indicating that overexpression of CFIm25 provided resistance to chemical inhibition of the NF-κB pathway. In conclusion, our data indicates that CFIm25 activates the NF-κB pathway and when overexpressed, it causes differentiation to happen more promptly than with PMA treatment alone.

**Fig 9.**
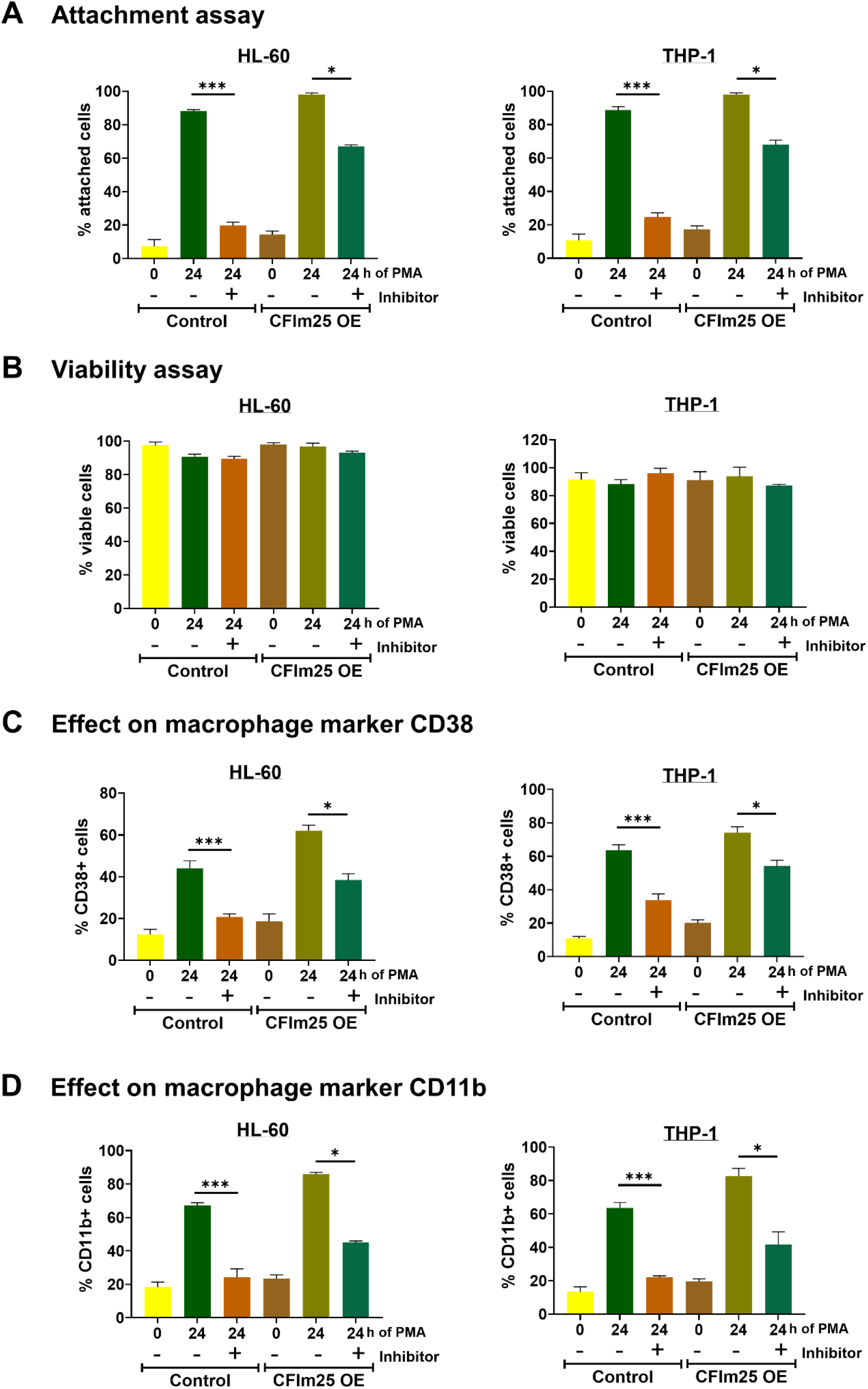
CFIm25 overexpression provides resistance to NF-κB inhibition. **(A)** Attachment assay. HL-60 and THP-1 control cells or cells overexpressing CFIm25 were incubated with or without the NF-κB inhibitor BAY 11-7082, and then treated with PMA for 0 and 24 hours and the percentage of cells attached at each time point determined. (B) Viability assay. The viability of cells was determined by the resazurin conversion assay. (C & D) Effect of NF-κB inhibition on macrophage markers CD38 and CD11b by flow cytometry of HL-60 and THP-1 control cells or those overexpressing CFIm25 with or without treatment with NF-κB inhibitor. The expression of CD38 and CD11b over time is shown as a percentage of CD38+ or CD11b+ cells. Data is representative of at least three biological replicates and is plotted as mean ± SE. P value <0.05 was considered significant, where * = P ≤ 0.05; *** = P ≤ 0.001.

## 3 Discussion

Monocyte differentiation yields macrophages primed to respond to stimuli that elicit innate immune functions such as inflammation or resolution of tissue damage. The goal of this study was to increase our understanding of how regulation of mRNA polyadenylation contributes to efficient macrophage differentiation. We had previously observed that CFIm25, a protein well-characterized for its ability to influence APA in other cell types, increased during macrophage differentiation of the THP-1 and U937 monocytic cell lines (6). In the current study, we have shown for the first time that this increase in CFIm25 has functional significance and facilitates multiple facets of the transition from monocytes to macrophages. We have found that CFIm25 promotes the cell cycle arrest that is a general characteristic of differentiating cells, the adherence to surfaces that allows cell migration, and the acquisition of the macrophage markers CD38 and CD11b.

Based on our findings, CFIm25 impacts macrophage differentiation in several ways, one of which is by activating the NF-κB pathway (Fig. 10). We chose the NF-kβ pathway to examine in greater detail because it is one of the major and best-documented pathways known to affect macrophage differentiation and function, and our differentiation inducer, PMA, is known to activate it (24, 46, 47). In our present study, several outputs of the NF-kβ pathway (27, 45) were enhanced by CFIm25 overexpression, including the Cyclin Dependent Kinase inhibitor p21, the anti-apoptotic and antiproliferative protein Bcl-XL, the adhesion molecule ICAM1, and the cytokine TNF-α. The increased phosphorylation of the p65 subunit of the NF-κB transcription factor and higher levels of the p50 subunit also indicate NF-κB activation. The CFIm25-mediated increase in TNF-α could further stimulate NF-κB activity (24) and production of CD38, a cell surface receptor characteristic of macrophages (48, 49). In addition, overexpression of CFIm25 provided resistance to inhibition of differentiation by the NF-κB inhibitor BAY 11-7082.

**Fig 10.**
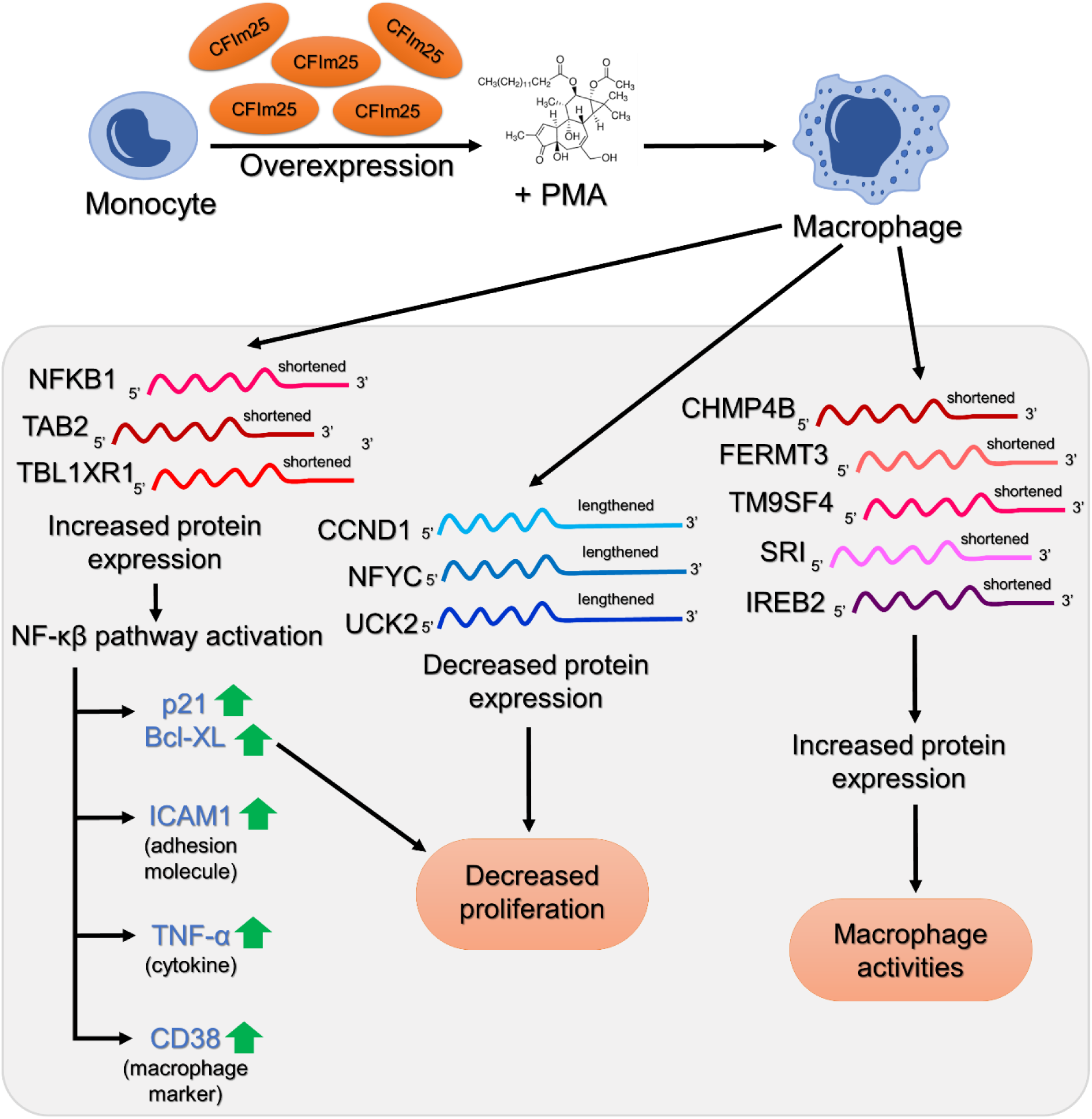
Summary of findings and a model to explain CFIm25-mediated regulation. Increased expression of CFIm25 in monocytic cells leads to more efficient differentiation to macrophages through regulated APA of critical genes and activation of the NF-κB pathway and its target genes.

APA of the mRNAs encoding effectors of the NF-κB pathway may contribute to the activation that we have observed. The 3’ UTRs of mRNAs of TAB2, TBL1XR1, and NFKB1 mRNAs are shortened, and there is a corresponding increase in the levels of both their mRNAs and proteins when CFIm25 is overexpressed. This is in accordance with the findings of Jia, et al (44) which documented the shortening of NFKB1 in virus-infected macrophages and showed that a luciferase reporter with the short NFKB1 3’ UTR made more protein than one with the long 3’ UTR. NFKB1 encodes the p105 precursor to the p50 subunit of NF-κB (50). TAB2 is part of the TAK1 kinase complex which is a driver of macrophage differentiation and is also needed for NF-κB activation during the inflammatory response to pathogens (28)(51, 52). TBL1XR1 binds the p65 subunit of NF-κB and promotes its nuclear translocation and transcription of NF-κB gene targets (29–31, 53).

It is interesting that the CFIm25-mediated increases in protein levels do not exactly track with increases in mRNA levels (Fig. 6 & 7). A possible explanation for this observation may be that changes in the 3’ UTR length can not only lead to changes in mRNA stability but also alter the translation efficiency of the mRNA without changing mRNA levels. It can also change the localization of the mRNA, such as whether it is exported from the nucleus (5) or associated with condensates that regulate translational activity (54). Changes in protein stability could also be a factor. Given the dramatic changes happening to the cell architecture, cell cycle progression, and the gene expression program during the 24-hour differentiation time course, it would not be surprising that the cells would use multiple mechanisms to achieve the optimum level of specific proteins in different phases of the differentiation process. While we do not know the exact underlying mechanisms, our study nevertheless demonstrates that CFIm25 overexpression increases the level of and/or hastens the expression of the NF-kβ p50, TAB2 and TBL1XR1 proteins during macrophage differentiation. We also note that the effects of CFIm25 during macrophage differentiation are likely not limited to the NF-κB pathway. NF-kβ signaling is closely coordinated with other signaling pathways such as PI3K/AKT, MAPK, JAK-STAT, TGF-β, Wnt, Notch, and Hedgehog (55). In fact, TBL1XR1 is thought to modulate some of these pathways in cancer cells (56). TAB2 activates MAPKs as well as NF-kβ (28, 57–59), both of which are necessary for the transition from monocyte to macrophages (60–62). CFIm25-mediated alternative polyadenylation may be one of the ways to modulate this crosstalk.

In the immune system, NF-κB is best studied for its role in inflammatory signaling (24, 45), but in many cellular contexts, NF-κB has a positive effect on cell cycle regulation through its ability to stimulate CCND1 (cyclin D1) transcription (27). However, several studies have shown that NF-κB can also induce cell cycle arrest at the G1/S transition in pro-B-cells and HeLa cells, and in epithelial cells and lymphoma cells, this arrest is attributable to NF-κB-induced transcription of CDKN1A (p21) (63–66). Delaying transition through the G1-S phase of the cell cycle is thought to be one of the hallmark events for the monocyte-macrophage transition (67), but a role for NF-κB has not been previously demonstrated. Upon overexpression of CFIm25, this delay is seen at an earlier point after induction of differentiation and at a later point when CFIm25 is depleted. Our findings suggest that a predominant consequence of the CFIm25-mediated NF-κB activation in differentiating monocytes is slowing of the G1-S phase, in part by increasing levels of the p21 and Bcl-XL.

Our findings indicate that CFIm25 can affect the cell cycle in ways that may not be dependent on the NF-κB pathway (Fig. 10). For example, CFIm25 has been shown to bind to UGUA elements in the 3’ UTR of CCND1 mRNAs and promote use of a distal PAS, leading to a decrease in Cyclin D1 protein and slowing of the G1-S phase. We observed this 3’ end lengthening and depletion of Cyclin D1 in monocytes stimulated to differentiate, and these changes were enhanced by CFIm25 overexpression. In addition, the mRNA 3’ end processing and protein levels of other potential regulators of cell cycle not characterized as NF-κB targets are altered after manipulation of CFIm25. For example, NFYC, a subunit of the CCAAT-binding transcription factor, drives the transcription of many genes needed for cell cycle progression, including the CFIm25-suppressed DNA replication protein PCNA (68), and inhibits p21 expression in glioma cells (69). UCK2 metabolically stimulates proliferation through its activity as a rate-limiting enzyme of the pyrimidine salvage pathway and non-metabolically by activating the EGFR-AKT pathway in hepatocellular carcinoma cells (70). CFIm25 overexpression causes lengthening of the mRNAs of NFYC and UCK2 and a corresponding decrease in mRNA and protein levels. In contrast, OAZ1 suppresses proliferation by promoting degradation of ornithine decarboxylase and Cyclin D1 (71), and has been proposed to induce erythroid differentiation (72). CFIm25 promotes accumulation of short OAZ1 mRNA isoforms and increased mRNA and protein. These findings support a multi-faceted regulation of cell cycle progression by CFIm25 during macrophage differentiation.

The APA elicited by CFIm25 on mRNA transcripts not known to be affected by NF-kβ signaling, such as those of CHMP4B/ESCRT-III, FERMT3, SRI, TM9SF4, and IREB2 may impact other critical macrophage activities (Fig. 10). The mRNAs of these genes are shortened and increased in level by CFIm25 overexpression, and this is associated with greater amounts of the encoded proteins. CHMP4B/ ESCRT-III is a component of the ESCRT machinery, which is involved in phagosome maturation and MHC-II antigen presentation (73). FERMT3, which encodes the integrin binding partner Kindlin-3, is needed for macrophage adhesion, migration and phagocytosis (74). TM9SF4 is required for cell adhesion and phagocytosis of Drosophila macrophages (75), and it may have a similar function in human macrophages. SRI (Sorcin) has not been connected to macrophage activity but is known to promote the migration of cancer cells (76), suggestive of a comparable role in macrophages. Another important function of macrophages is to recycle iron obtained from phagocytosis of old or damaged red blood cells (77). IREB2 (Iron-Responsive Element-Binding Protein 2) facilitates this by regulating the translation and stability of mRNAs for the iron storage protein ferritin. Together, these associations indicate a significant role for CFIm25 in the production of functional macrophages and point to possible uncharacterized modulators of macrophage differentiation and function.

Regarding mechanism, CFIm25 is probably collaborating with other APA regulators to generate the optimal array of mRNA isoforms. For example, we and others have documented a role for the CstF64 C/P protein in promoting macrophage differentiation (6) and the inflammatory response to lipopolysaccharide (78), and RNA-binding protein SRSF12 regulates APA events in ways that likely contribute to a pro-inflammatory phenotype (79). Depletion of FIP1L1, one of the core cleavage and polyadenylation proteins that also regulate APA, promoted a more differentiated state in Acute Myeloid Leukemia cells as evident from reduced cell surface expression of the hematopoietic stem cell marker CD34, a decrease in cell proliferation, and an increase in mRNAs of classical markers of mature myeloid cells (80). Numerous proteins have now been identified as APA regulators (3–5) and knowledge of how they might influence macrophage differentiation and interact with each other will be important to understanding the full impact of APA on this transition. CFIm25 can also promote RNA Polymerase II transcription in a way that is independent of its function in 3’ end processing (81), and it can influence alternative splicing events and the generation of circular RNAs that can compete for miRNAs (14). However, it is not known how these CFIm25 targets impacts macrophage differentiation.

In conclusion, we show that CFIm25 expedites the process of differentiation from monocytes to macrophages. Our findings support a model in which CFIm25 accomplishes this task by mediating alternative polyadenylation of key genes, including ones involved in cell cycle regulation and NF-κB signaling. Future research might lead to ways to manipulate the polyadenylation machinery or the APA of specific transcripts to better promote the resolution of infections, cancer, and other diseases affected by macrophage activity.

## 4 Materials and methods

### 4.1 Cell culture, treatment, and microscopy

All cells were maintained in RPMI 1640 medium supplemented with 2 mM L-glutamine and 10% heat-inactivated fetal bovine serum (FBS) at 37°C in a humidified atmosphere of 5% CO_2_. Differentiation of the HL-60 (ATCC® CCL-240) and THP-1 (ATCC® TIB-202) human monocytic cells into macrophages was performed by treatment with 3 nM phorbol-12-myristate-13-acetate (PMA; Sigma-Aldrich) for up to 24 hours. Cells were cultured in 6-well plates at a density of 1×10^6^ cells/well. For the NF-κB inhibitor assays, HL-60 and THP-1 cells in both control and overexpressing CFIm25 groups were treated with 5 µM BAY-11-7082, and after 24 hours, the cells were treated with PMA, followed by subsequent analyses.

### 4.2 Attachment, proliferation and viability assays

HL-60 and THP-1 cells were either untreated or treated with PMA and left for the indicated times. After that, unbound cells were collected from the plate by washing gently with phosphate-buffered saline (PBS) and collected for counting (denoted as “suspended”). The attached cells were collected by scraping to release them from the plate. Cells were then stained with trypan blue and counted in a hemocytometer and represented as percentage of suspended or attached cells over total (suspended + attached). The figures are an average of three independent observations. The number of live cells before and after differentiation were calculated by trypan blue exclusion to see how well the cells proliferated in each condition. For the viability assay, we used resazurin for a fluorescent assay that detects cellular metabolic activity. The blue nonfluorescent resazurin reagent is reduced to highly fluorescent resorufin by dehydrogenase enzymes in metabolically active cells. This conversion only occurs in viable cells and thus, the amount of resorufin produced is proportional to the number of viable cells in the sample. The resorufin formed in the assay is quantified by measuring the relative fluorescence units (RFU) at 590-620 nm.

### 4.3 Cell cycle analysis by DNA content

HL-60 and THP-1 cells were seeded in fresh RPMI media with 10% FBS for 24 h before cell cycle analysis. After PMA treatment, cells were harvested at different time points (0-24h) with trypsinization and fixed in 70% ethanol overnight at −20°C. After washing with PBS, the cells were incubated with DNase-free RNase A (Sigma) at 5 ug/ml final concentration and incubated at 37°C for 30 min. Cell cycle was studied by PI (propidium iodide) staining using the Fluorescence-Activated Cell Sorter (FACS verse, with software FACSuite, V1.0.5.3841 Becton Dickinson, USA). A total of 10,000 events were acquired for each sample. Two types of gates were applied. First, the cells were gated according to their size and granularity to exclude debris, and then doublet discrimination was performed by gating the cells by area vs width to identify the singlet population. DNA content was analyzed for this gated population to give a histogram of DNA content (X-axis, PI fluorescence) versus counts (Y-axis).

### 4.4 Western blot analysis

Western blots were performed with total cell extracts of untreated and differentiated cells. For preparation of extract, RIPA buffer (20 mM Tris-HCl pH 7.5, 150 mM NaCl, 1 mM Na_2_EDTA, 1 mM EGTA, 1% NP-40, 1% sodium deoxycholate, 2.5 mM sodium pyrophosphate, 1 mM β-glycerophosphate, 1 mM Na_3_VO_4_ and 1 µg/ml leupeptin) was added to the cell pellet, followed by incubation on ice for 15 min, centrifugation at 25,000 rcf for 10 mins at 4°C, and supernatant collection. Protein concentration was measured with the BCA reagent (Pierce, Thermo Fisher Scientific), and 50-80 µg protein was separated on a 10% polyacrylamide-SDS gel and transferred to PVDF membrane. Ponceau Red staining was done for total protein staining of blots. The membrane was then cut into segments to enable probing for multiple proteins from the same blot. For overexpression and knockdown experiments, control and treated samples were run on the same gel. The membrane was then blocked for 1 h in 5% non-fat dried milk in TBS-T (Tris Buffer Saline with Tween-20, i.e., 0.05% Tween 20 in 1X TBS) buffer followed by rocking overnight at 4°C with primary antibodies (antibody sources provided in Supplementary Table 1). Unbound antibodies were removed by 4 × 5 min washes with TBS-T buffer. The membrane was then incubated with an HRP-conjugated secondary antibody at a dilution of 1:5000 for 1 h at room temperature followed by 3 × 5-min washes with TBS-T and a final wash with TBS. The blot was developed with SuperSignal™ West Pico PLUS or Femto Chemiluminescent Substrate (Thermo Fisher Scientific), visualized with a ChemiDoc XRS+ System (Bio-Rad), and quantitated using Image J software (82). GAPDH, β-actin and Lamin B1 were used as control for individual western blots because their levels did not change upon differentiation. Pre-stained protein markers were used as internal molecular mass standards, and each western blot was performed in three biological replicates of each time point of the differentiation process.

### 4.5 Flow cytometry analysis for macrophage markers

Surface marker analysis of macrophages was performed using flow cytometry. After differentiation, cells were recovered from culture plates by gentle scraping. Cells were washed in 1X Phosphate Buffered Saline (containing 137 mM NaCl, 2.7 mM KCl, 8 mM Na2HPO4, and 2 mM KH2PO4), 1% BSA, 0.01M NaN3 solution and incubated with Human TruStain FcX™ (Fc blocker, Biolegend) for 15 min to block Fc receptors and reduce nonspecific binding. Thereafter, cells were stained with 1:20 of PE-tagged CD38 or PE-tagged CD11b (macrophage marker) antibodies for 30 min on ice. Cells were fixed with buffer (Cytofix, Becton-Dickinson Biosciences), according to the manufacturer’s protocol. Samples were acquired on a BD LSRII flow cytometer, and data were analyzed using FACS diva or FlowJo. For analysis, samples were gated on light scattering properties to exclude dead cells and debris. Unstained control samples were used to determine the level of background fluorescence. These controls help in setting the baseline for autofluorescence in the cells, ensuring that the PE signal detected is specifically due to the antibody staining. Fluorescence-minus-one (FMO) controls were used to set accurate gates for distinguishing between positive and negative populations. FMO controls include all the fluorochromes used in the experiment except the one being measured, allowing for the assessment of spillover and background fluorescence. This helps to precisely define the boundary between PE-positive and PE-negative cells, improving the accuracy of the gating strategy. Here, we have used APC, Brilliant violet (BV711) or FITC as the FMO control.

### 4.6 RNA sequencing by QuantSeq

Library preparation of total RNA was carried out by using the Lexogen 3′ mRNA-Seq QuantSeq FWD kit according to manufacturer’s instructions. Library preparation, quality control, and sequencing were carried out by Admera Health (South Plainfield, NJ, USA). cDNA libraries were sequenced on an Illumina HiSeq machine (2 × 150 nt) at Admera Health.

### 4.7 APA analysis

The reverse read (read 2) of QuantSeq FWD was used for APA analysis. Briefly, adapter and poly(T) regions were trimmed and trimmed reads were aligned by using STAR-2.7.7a to the human genome sequence (hg19). Only the alignments to PAS regions annotated by the PolyA_DB v3 database (−100 to +25 nt around each PAS) were kept. The last aligned position (LAP) of each mapped read was compared to PolyA_DB-annotated-PASs (83), allowing ± 24 nt flexibility. Matched reads were considered PAS-supporting (PASS) reads, which were used for further APA analysis. The abundance of transcripts associated with a given PAS was based on the number of PASS reads normalized to the total number of PASS reads per sample, yielding a reads per million (RPM) value. UCSC genome browser tracks were generated by bedtools v2.31.0 (84) and bedGraphToBigWig (85).

For 3′ UTR APA analysis, we used DEXSeq analysis (86), where the two PASs with the highest usage levels in the 3′ UTR of the last exon were compared. One was named proximal PAS (pPAS) isoform, and the other distal PAS (dPAS) isoform. The difference in log_2_[reads per million (RPM) ratios] of the two PAS isoforms between two samples (control and CFIm25 overexpression) was defined as relative expression difference (RED). Significant APA events were those with RED > log_2_(1.2) or < −log_2_(1.2) and p < 0.05.

### 4.8 RT-qPCR analysis

Total RNA extraction was carried out on 2×10^6^ cells using Trizol according to manufacturer’s protocol. 1.5 µg of RNA was subjected to reverse transcription using oligo dT primer and Superscript III reverse transcriptase. The cDNA was amplified by qPCR with specific primers (Supplementary Table 2) using the C1000™ thermal cycler with CFX96 Touch Real-Time PCR Detection System (Bio-Rad Laboratories). The relative expression of genes was analyzed quantitatively by the ΔΔC_t_ method. ACTB RNA was used as the normalization control for RT-qPCR-based RNA expression analyses because its level did not change upon differentiation according to the QuantSeq datasets. Primers for total or long transcripts were designed according to the PAS annotation featured in the PolyA_DB database and the PAS that was affected by PMA treatment (as observed in the UCSC genome browser representation of the QuantSeq data).

### 4.9 Lentivirus construction, siRNA and cell treatment

The OE-CFIm25 lentivirus vector was a kind gift from Shervin Assassi, University of Texas Health Science Center at Houston, TX (23). It was generated by cloning the coding sequence (CDS) of human CFIm25 into the pLV-EF1a-IRES-Puro Vector (Addgene) that contains an EF-1a promoter upstream of an IRES element to co-express the puromycin marker. The CFIm25 CDS was inserted between the EF-1a promoter and IRES, which allows the expression of CFIm25 and the puromycin marker from a single mRNA sequence. The vector backbone without an insert was used as OE-control. For CFIm25 KD, ON-TARGETplus SMARTpool human siRNAs against CFIm25 were purchased from Horizon Discovery Biosciences Limited.

### 4.10 Lentiviral transfection

The recombinant plasmids were co-transfected with the components of Dharmacon™ Trans-Lentiviral packaging kit into HEK293FT cells using FuGENE® HD (Promega Corp.) according to manufacturer’s protocol. Transfection of HEK293FT cells was carried out in 6-well plates when the cells were 80-85% confluent, and transfection media were changed after 16 h. The recombinant lentiviruses were harvested at 48 h post-transfection, spun at 1250 rpm for 5 mins and filtered by a 0.45 µm filter to remove cells and debris. Purified viruses were used for infecting monocytic cells. 1×10^6^ target cells were seeded in 2 mL of media per well in a 6-well plate and cultured overnight. Lentiviral particles (2 mL per well) were added the next day to the cells in culture media containing 10 µg/mL polybrene for efficient infection. Selection of cells stably expressing OE-control and OE-CFIm25 started 72 h post-transfection. Growth medium was replaced with fresh selection medium containing 1 μg/mL of puromycin. Puromycin-containing medium was refreshed every 2–3 days, and selection was completed after approximately 1 week, after which clones were expanded for 2 more weeks and then frozen for later use.

### 4.11 siRNA transfection

For CFIm25 KD, cells are seeded, allowed to rest for 1 hour, then transfected with 50 nM of siRNA pool with jetOPTIMUS® transfection reagent (Polyplus). This was followed by a 6-hour resting time for the cells to take in RNA and recover. Cells are then distributed evenly into six plates that are used for the different time points. Cells are harvested for the 0 hour and PMA is added to the remaining plates for the other time points. The time gap between seeding of transfected cells and PMA treatment is 7 hours.

### 4.12 ELISA for TNF-α

The BioLegend® Legend Max™ kit was used to perform ELISA according to manufacturer’s protocol. In this sandwich ELISA, a human TNF-α specific monoclonal antibody is precoated on a 96-well strip-well plate. Supernatants from PMA-treated HL-60 and THP-1 cells are used to quantify the amount of expressed TNF-α, which was represented as pg/mL.

### 4.13 Statistical analysis

All experiments were performed in at least three independent sets. Data are presented as mean ± SE. Statistical analysis was performed using GraphPad Prism 6.01 (GraphPad Software Inc., La Jolla, CA, USA). Two-way analysis of variance (ANOVA) was performed to determine the significance between the groups. Considerations were * = P ≤ 0.05; ** = P ≤ 0.01; *** = P ≤ 0.001. A p value <0.05 was considered significant.

## 5 Conflict of Interest

The authors have no relevant financial or non-financial interests to disclose.

## 6 Author Contributions

CM, SM and AB conceived the study and CM provided general oversight. SM and AB developed the strategy and methodology, designed the experiments, acquired the data, reported and organized the findings. All authors interpreted the results. The first draft of the manuscript was written by SM and all authors commented on previous versions of the manuscript. LW and BT contributed to the computational analyses. All authors contributed to the article and approved the submitted version.

## 7 Funding

This work was supported by the National Institutes of Health 1R01AI152337 to CM, R01GM084089 to BT, and the Natalie Zucker Research Award to SM.

## Abbreviations

C/P: cleavage and polyadenylation
PMA: Phorbol Myristate Acetate
APA: Alternative polyadenylation
PAS: Poly(A) Site

## 9 Data Availability Statement

The datasets generated during and/or analyzed during the current study are available in the GEO repository with accession number GSE254061.

## Notes

### Competing Interest Statement

The authors have declared no competing interest.

