## Supplementary information for "The alternative polyadenylation regulator CFIm25 promotes macrophage differentiation and activates the NF-κβ pathway"

*Supplementary Material*

### 1 Supplementary Tables and Figures

#### 1.1 Supplementary Tables

**Supplementary Table 1. Antibodies used in this study:**

**Western blot antibodies:**

| Catalogue no. | Name | Host | MW | Vendors |
|---|---|---|---|---|
| SC-81109 | CFIm25 | Mouse monoclonal | 26 kDa | Santa Cruz Biotechnology |
| SC-374650 | CD38 | Mouse monoclonal | 45 kDa | Santa Cruz Biotechnology |
| SC-8396 | Cyclin D1 | Mouse monoclonal | 37 kDa | Santa Cruz Biotechnology |
| SC- 56 | PCNA | Mouse monoclonal | 36 kDa | Santa Cruz Biotechnology |
| SC-47778 | β-actin | Mouse monoclonal | 43 kDa | Santa Cruz Biotechnology |
| SC-365062 | GAPDH | Mouse monoclonal | 37 kDa | Santa Cruz Biotechnology |
| SC-374015 | Lamin B1 | Mouse monoclonal | 67 kDa | Santa Cruz Biotechnology |
| 3033S | Phospho NF-κβ-p65 | Rabbit monoclonal | 65 kDa | Cell Signaling Technologies |
| 8242S | NF-κβ-p65 | Rabbit monoclonal | 65 kDa | Cell Signaling Technologies |
| SC-8414 | NF-κβ-p50 | Mouse monoclonal | 50 kDa | Santa Cruz Biotechnology |
| SC-8439 | ICAM1 | Mouse monoclonal | 85-110 kDa | Santa Cruz Biotechnology |
| SC-6246 | p21 | Mouse monoclonal | 21 kDa | Santa Cruz Biotechnology |
| SC-8392 | Bcl-XL | Mouse monoclonal | 30 kDa | Santa Cruz Biotechnology |
| SC-398188 | TAB2 | Mouse monoclonal | 83 kDa | Santa Cruz Biotechnology |
| SC-100908 | TBL1XR1 | Mouse monoclonal | 55 kDa | Santa Cruz Biotechnology |
| 28329-1-AP | IREB2 | Rabbit polyclonal | 95-110 kDa | Proteintech |
| Ab-7839 | FERMT3 | Rabbit polyclonal | 76 kDa | Abclonal Antibody |
| 136183-1-AP | CHMP4B | Rabbit polyclonal | 25-33 kDa | Proteintech |
| 25595-1-AP | TM9SF2 | Rabbit polyclonal | 70 kDa | Proteintech |
| SC-100859 | SORCIN | Mouse monoclonal | 22 kDa | Santa Cruz Biotechnology |
| A7444 | OAZ1 | Rabbit polyclonal | 25 kDa | Abclonal Antibody |
| SC-390985 | NFYC | Mouse monoclonal | 40 kDa | Santa Cruz Biotechnology |
| 422301 | Human TruStain FcX™ | Monoclonal Antibody | $F_c$ blocker | Biolegend |

**Flow cytometry antibody:**

| Catalogue no. | Name | Host | Fluorophore |
|---|---|---|---|
| Biolegend 102707 | CD38 | Mouse | PE |
| Proteintech PE-65116 | CD11b | Mouse | PE |

**Supplementary Table 2. Primers used in this study:**

| Purpose | Primer name | Forward primer 5'-3' | Reverse primer 5'-3' |
|---|---|---|---|
| RT-qPCR for gene expression | ACTB total | CATGTACGTTGCTATCCAGGC | CTCCTTAATGTCACGCACGAT |
| | CCND1 total | AACTACCTGGACCGCTTCCT | CCACTTGAGCTTGTTCACCA |
| | NFYC total | GGACTCCTGAGCAGAGTTGT | GCAAAGAGTACAGGCGCTTC |
| | UCK2 total | CGGAAAGCGGAGGGAGTC | CTATAAGGAAGGGCTCGCC |
| | CHMP4B total | GGTGTTCGGGAAGCTGTTC | TTTTGGTGCCGTGCTTCTTG |
| | FERMT3 total | AGAGAAAGGAGGGCAGGAAG | TCCTCCTCTCCCACAAACAC |
| | IREB2 total | AGCCTGCTTCCTTCTTTCCT | CCATCGCCGGAGACCATATT |
| | OAZ1 total | GCATCTATAAAGGCGGGCG | CCTTCCTTCTCTCTGGCGAA |
| | SRI total | CTCAGGATCCGCTGTATGGT | TGTATCCTCCAGCAATGCCA |
| | TAB2 total | AGGGAGGGCTGAGGTGTC | CACTAGGGCAGCCGTCTC |
| | TBL1XR1 total | CTGTCCCTGATAGATGCCGT | TTCTCCTCCCCATTTGCTGT |
| | TM9SF2 total | CAACTATCATGAGCGCGAGG | CTTGCACTCGTCGCTCTTTT |
| | NFKB1 total | CAAGCAGCTCTGCAGCAG | ACTGTCATAGATGGCGTCTG |
| RT-qPCR of long transcript for APA analysis | CCND1 long | ACGCTTTGTCTGTCGTGATG | GTGCAACCAGAAATGCACAG |
| | NFYC long | CCAAGACTTGCCACGTTGTT | GGCAATGAATCCACCCACTC |
| | UCK2 long | GCCTCTCACTCCTTCACACT | CAAGGTGAACTGGAGACCCT |
| | CHMP4B long | CTTGCCGCACATCTCTTTGT | AAACTCCAGAAGACCAGGGG |
| | FERMT3 long | GGCCAGACGCTGTACC | CAAGAAAGAACTCGTTTGAAAC |
| | IREB2 long | GGTAACAAGGTCGTGTGCAT | AAGCCAAAGTCCACCCTCTT |
| | OAZ1 long | GTGTTTGTGATACTGAAGTATTT GC | CAAGTTAAAAGACTAAGACTGTTT CC |
| | SRI long | TGAGCCATTATCAGTCATGCC | ACCCAAGTGCGTCTATGTCA |
| | TAB2 long | AGACCCAAAGCCCTTACGTT | ACCCAGCTAAATCACAAAACCA |
| | TBL1XR1 long | TAAACCAGCCCATGACAGGT | AGGGGAACTGCAACAACAAC |
| | TM9SF2 long | CCTGGACATTAGCAATCACTAG C | CAGCAAGCAGAGAGACCCTA |
| | NFKB1 long | CGTTCCTATTGTCATTAAAGGTA TC | ATGGCACATCAAGTGACTCTC |

**Supplementary Table 3. Outputs for APA events identified by 3' Quant sequencing.** Table outlining the genes undergoing shortening or lengthening on CFIm25 overexpression from poly(A) Quant-sequencing. In this table chr: chromosome number where the gene is located, strand: the strand of chromosome on which the gene is coded, Prx_pA_pos: position of the proximal poly(A) site, Dis_pA pos: position of the distal poly(A) site, RED: relative expression difference (RED) between proximal and distal p(A) isoforms in OE vs. control samples (RED = difference in $log_2$(ratio) of read numbers of two p(A) isoforms between two samples), pval.fisher.adj: p value calculated for the RED score by Fisher's exact test and change: whether the gene is significantly shortened or lengthened.

| Symbol | Description | chr | Strand | Prx_pA _pos | Dis_pA pos | RED | pval. fisher. adj | Change |
|---|---|---|---|---|---|---|---|---|
| **NFYC** | nuclear transcription factor Y subunit gamma | chr 1 | + | 41,236,778 | 41,237,273 | 1.39 | 2.07e-02 | lengthened |
| **UCK2** | uridine-cytidine kinase 2 | chr 1 | + | 165,877,345 | 165,880,855 | 0.99 | 1.99e-02 | lengthened |
| **CHMP4B** | charged multivesicular body protein 4B | chr 20 | + | 32,441,636 | 32,442,169 | -0.82 | 1.36e-02 | shortened |
| **FERMT3** | FERM domain containing kindlin 3 | chr 11 | + | 63,991,299 | 63,991,363 | -0.95 | 2.49e-02 | shortened |
| **IREB2** | iron responsive element binding protein 2 | chr 15 | + | 78,792,879 | 78,793,794 | -0.63 | 2.54e-02 | shortened |
| **OAZ1** | ornithine decarboxylase antizyme 1 | chr 19 | + | 2,273,238 | 2,273,323 | -0.87 | 1.36e-02 | shortened |
| **SRI** | sorcin | chr 7 | - | 87,835,606 | 87,834,431 | -1.1 | 1.01e-02 | shortened |
| **TAB2** | TGF-beta activated kinase 1 (MAP3K7) binding protein 2 | chr 6 | + | 149,732,659 | 149,732,741 | -1.26 | 2.90e-02 | shortened |
| **TBL1XR1** | TBL1X/Y related 1 | chr 3 | - | 176,741,135 | 176,738,543 | -0.85 | 3.75e-02 | shortened |
| **TM9SF2** | transmembrane 9 superfamily member 2 | chr 13 | + | 100,215,064 | 100,215,643 | -1.01 | 1.01e-02 | shortened |

**Supplementary Figures.**

**Supplementary Figure 1. Flow cytometry analysis of cells with and without overexpression of CFIm25. (A & B) Raw data for cell cycle analysis.** Representative raw data for cell cycle analysis of HL-60 (A) and THP-1 (B) cells treated with PMA for the indicated hours. Cell cycle was studied using PI staining as described in Materials and Methods, followed by flow cytometry. A total of 10,000 events were acquired for each sample. Two types of gates were applied. First, the cells were gated according to their size and granularity and next, doublet discrimination was performed by gating the cells as area vs width. DNA content was analyzed for this gated population. A histogram of DNA content (X-axis, PI fluorescence) versus cell counts (Y-axis) is displayed. **(C & D) Raw data for flow cytometry.** Representative dot plots for flow cytometry of macrophage marker CD38 at different time points during differentiation of HL-60 (C) and THP-1 (D) cells, where the percentage represents cells positive for CD38. Samples were prepared according to Materials and Methods and acquired on a BD LSRII flow cytometer, and data were analyzed using FACS diva. For analysis, samples were gated on light scattering properties to exclude dead cells and debris. Unstained control samples were used to determine the level of background fluorescence. Each dot plot displays FMO control on the x-axis and PE-tagged CD38 on the y-axis.

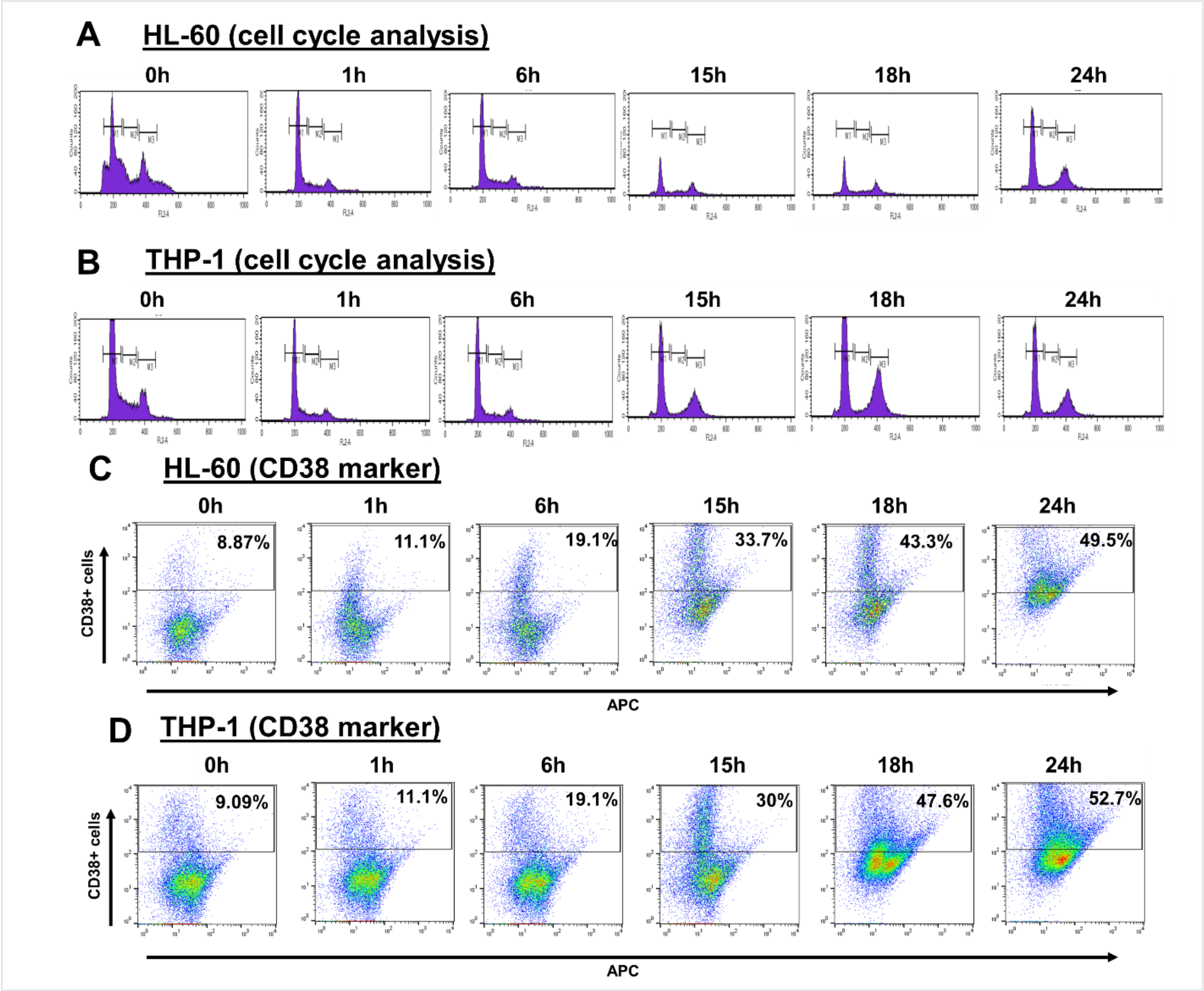

**Supplementary Fig 2. Effects of CFIm25 overexpression. (A & B) Attachment and viability assays in (A) HL-60 and (B) THP-1 cells overexpressing CFIm25.** For the attachment assay (*left*), cells overexpressing CFIm25 and control were treated with PMA for 0, 6 and 24 hours and live cells visualized and counted by the Trypan blue exclusion assay with respect to control. The graph presents the percentage of cells that are suspended or attached at each time point. For the viability assay (*right*), the viability of cells for OE-CFIm25 with respect to OE-control after differentiation indicates the amount of resazurin converted to resorufin produced, which is proportional to the number of viable cells in the sample, compared to those in the absence of PMA. The figure represents mean ± SE from three independent experiments. **(C & D) Raw data for cell cycle analysis.** Flow cytometry data of HL-60 (C) and THP-1 (D) cells overexpressing CFIm25 and treated with PMA for indicated hours with respect to control, where the y-axis represents cell numbers, and the x-axis represents incorporation of propidium iodide (PI). Gating is performed according to Supplementary Fig. 1A & B. Data is representative of at least three biological replicates. **(E-F) CD38 levels in genetically manipulated HL-60 (E) and THP-1 (F) cells by flow cytometry.** Flow cytometry staining of the surface marker CD38 at different time points during differentiation of control cells and cells overexpressing CFIm25, where percentage represents cells positive for CD38. Gating is performed according to Supplementary Fig. 1C & D. Data is representative of at least three biological replicates.

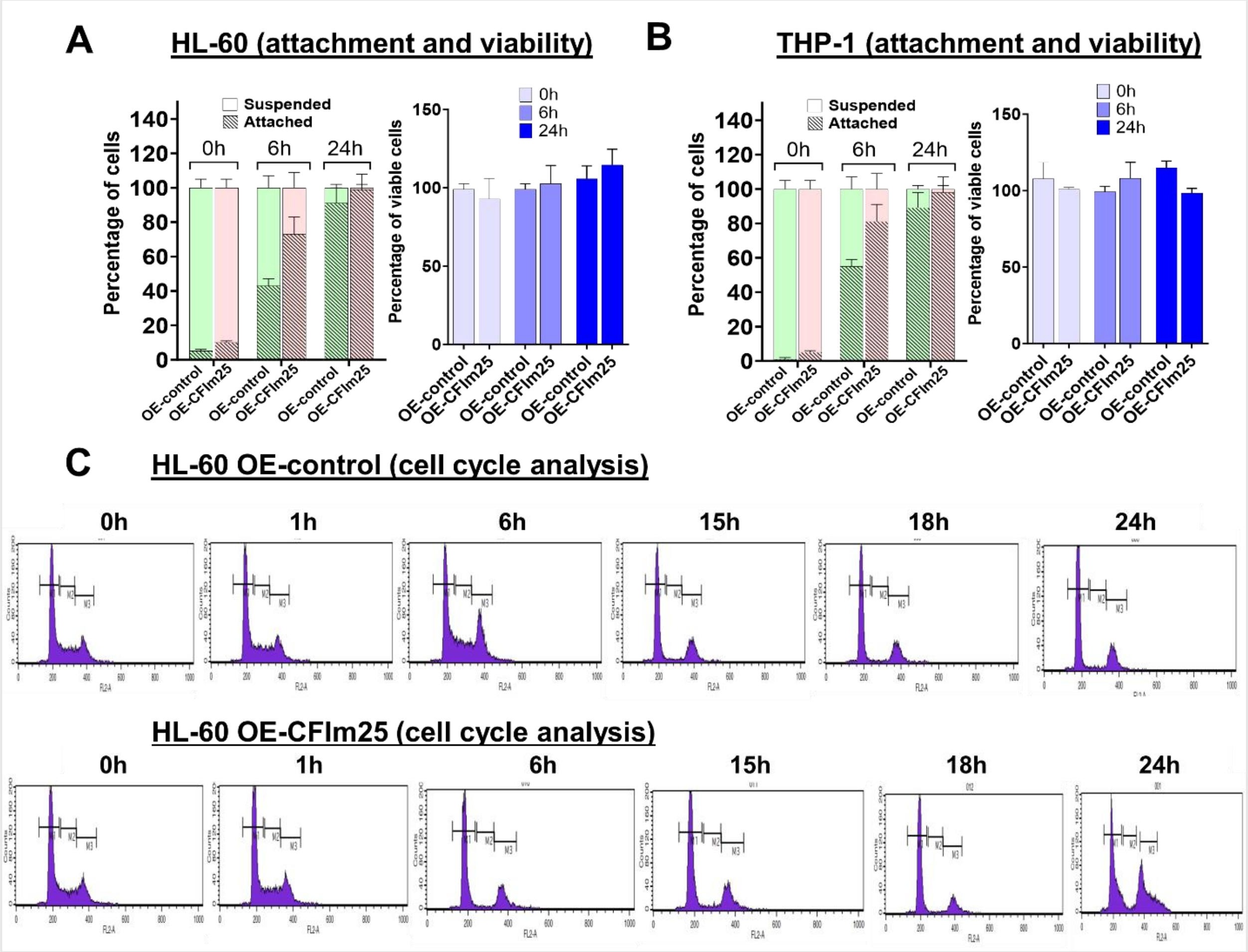

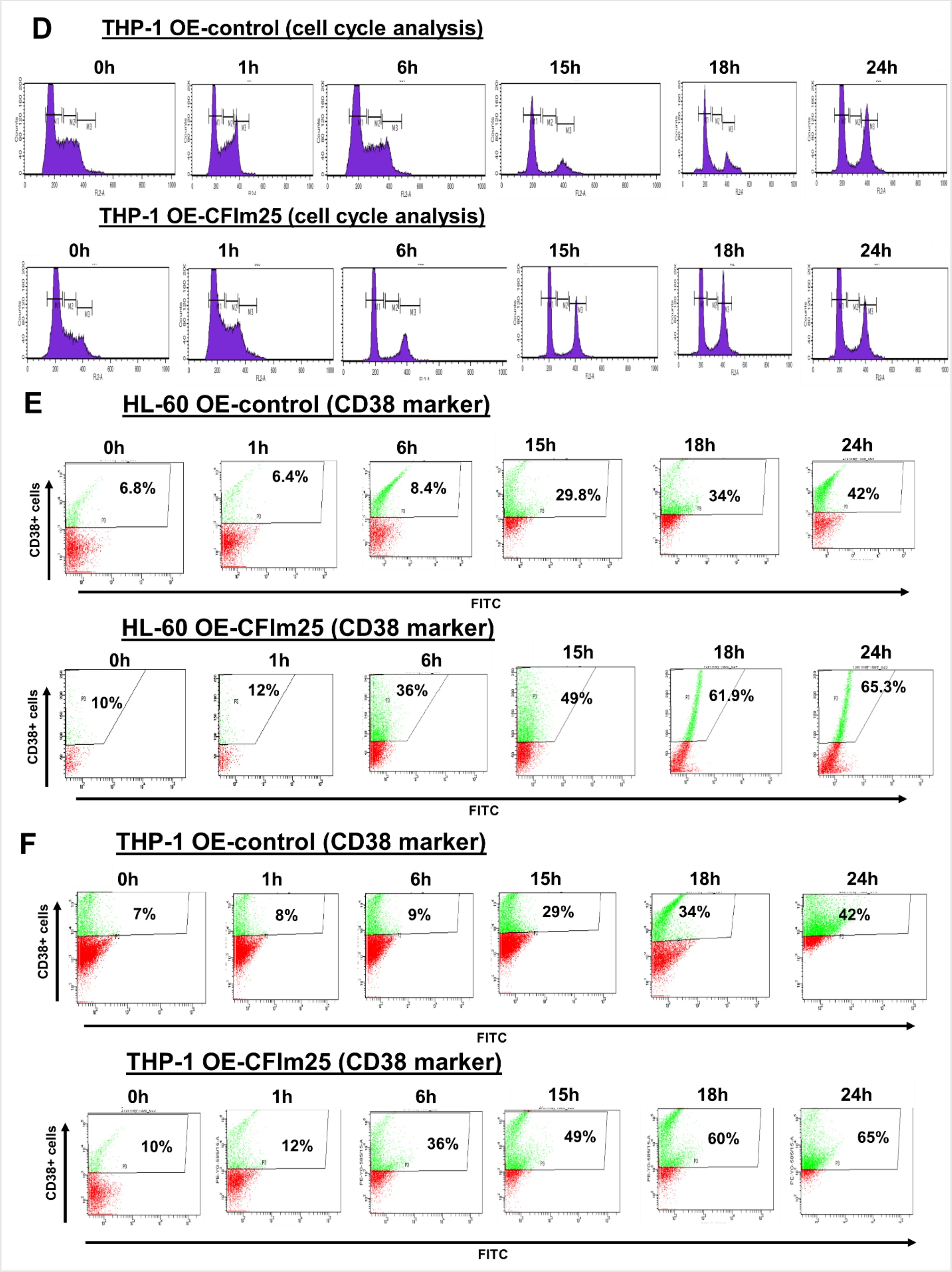
D
THP-1 OE-control (cell cycle analysis)
0h
1h
6h
15h
18h
24h
THP-1 OE-CFIm25 (cell cycle analysis)
0h
1h
6h
15h
18h
24h
E
HL-60 OE-control (CD38 marker)
CD38+ cells
0h 6.8%
1h 6.4%
6h 8.4%
15h 29.8%
18h 34%
24h 42%
FITC
HL-60 OE-CFIm25 (CD38 marker)
CD38+ cells
0h 10%
1h 12%
6h 36%
15h 49%
18h 61.9%
24h 65.3%
FITC
F
THP-1 OE-control (CD38 marker)
CD38+ cells
0h 7%
1h 8%
6h 9%
15h 29%
18h 34%
24h 42%
FITC
THP-1 OE-CFIm25 (CD38 marker)
CD38+ cells
0h 10%
1h 12%
6h 36%
15h 49%
18h 60%
24h 65%
FITC

**Supplementary Fig 3. Effect of CFIm25 knockdown. (A & B) Attachment and viability assays for (A) HL-60 and (B) THP-1 control cells or cells knocked down with siRNA against CFIm25.** For the attachment assay (*left*), cells treated with siRNA against CFIm25 and control were treated with PMA for 24 hours and live cells visualized and counted by the Trypan blue exclusion assay. The graph presents the percentage of cells that are suspended or attached. For the viability assay (*right*), the number of cells for si-CFIm25 with respect to si-control after differentiation indicates the amount of resazurin converted to resorufin produced, which is proportional to the number of viable cells in the sample, compared to those in the absence of PMA. The figure represents mean ± SE from three independent experiments. **(C & D) Raw data for cell cycle analysis.** Flow cytometry data of HL-60 (A) and THP-1 (B) cells knocked down for CFIm25 and treated with PMA for indicated hours with respect to control, where the y-axis represents cell numbers, and the x-axis represents incorporation of propidium iodide (PI). Gating is performed according to Supplementary Fig. 1A & B. Data is representative of at least three biological replicates. **(E-F) CD38 levels in genetically manipulated HL-60 (E) and THP-1 (F) cells by flow cytometry.** Flow cytometry staining of surface marker CD38 at different time points during differentiation of control cells and cells knocked down with siRNA against CFIm25, where percentage represents cells positive for CD38. Gating is performed according to Supplementary Fig. 1C & D. Data is representative of at least three biological replicates.

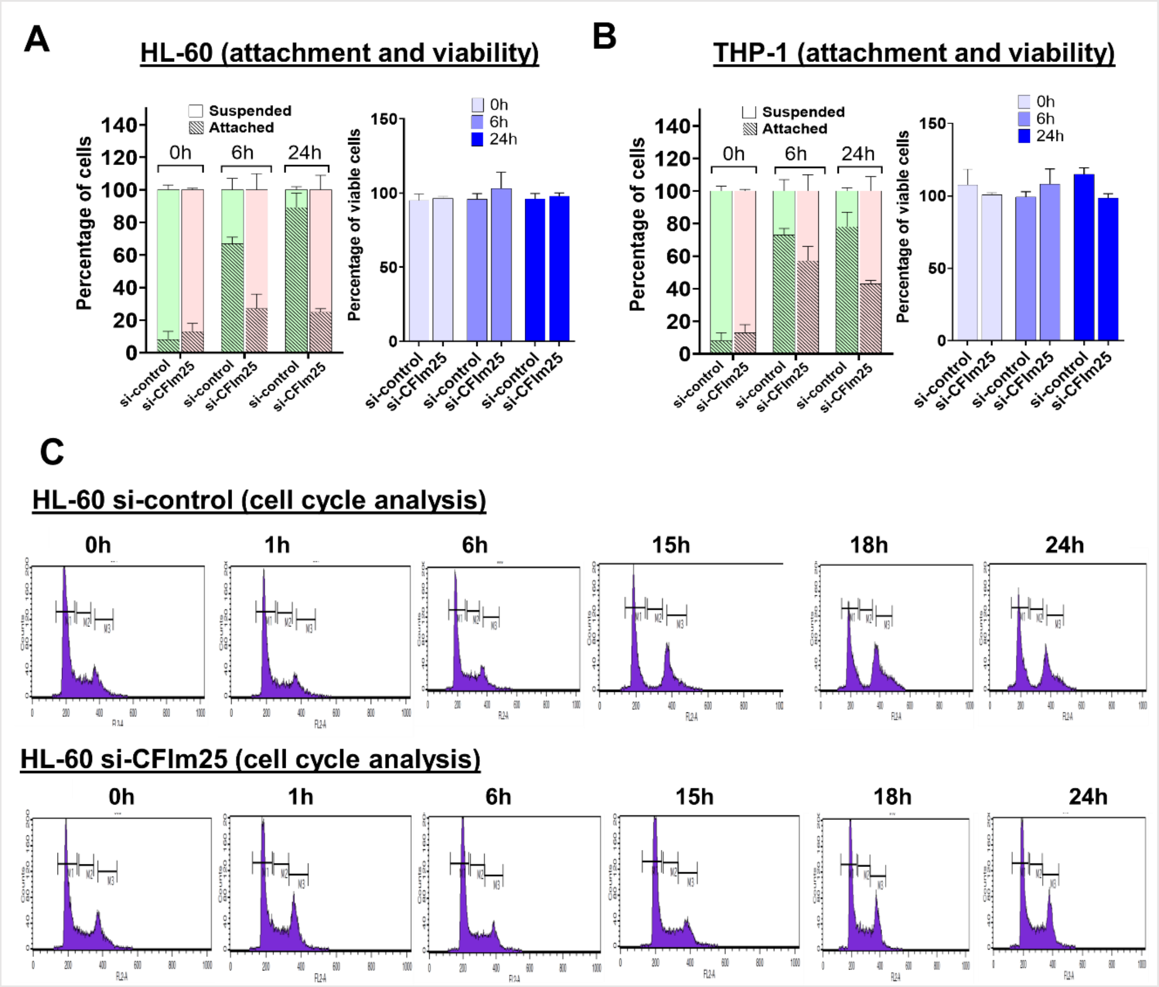

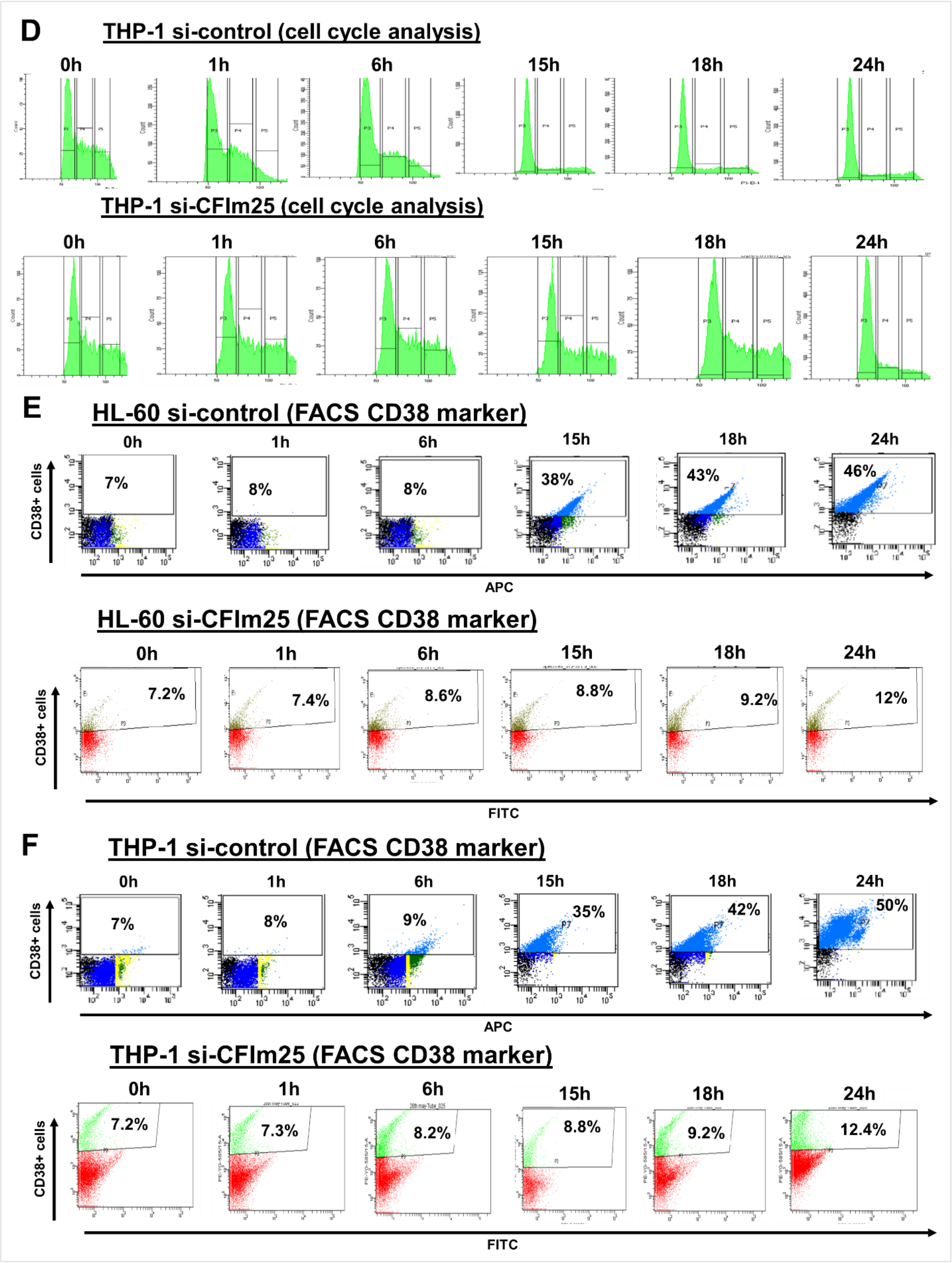
D
THP-1 si-control (cell cycle analysis)
0h
1h
6h
15h
18h
24h
THP-1 si-CFIm25 (cell cycle analysis)
0h
1h
6h
15h
18h
24h
E
HL-60 si-control (FACS CD38 marker)
0h
1h
6h
15h
18h
24h
CD38+ cells
7%
8%
8%
38%
43%
46%
APC
HL-60 si-CFIm25 (FACS CD38 marker)
0h
1h
6h
15h
18h
24h
CD38+ cells
7.2%
7.4%
8.6%
8.8%
9.2%
12%
FITC
F
THP-1 si-control (FACS CD38 marker)
0h
1h
6h
15h
18h
24h
CD38+ cells
7%
8%
9%
35%
42%
50%
APC
THP-1 si-CFIm25 (FACS CD38 marker)
0h
1h
6h
15h
18h
24h
CD38+ cells
7.2%
7.3%
8.2%
8.8%
9.2%
12.4%
FITC

**Supplementary Fig 4. Changes in poly(A) site usage after 6 hours of PMA treatment in control HL-60 cells compared to those overexpressing CFIm25. (A & B)** UCSC genome browser plots of merged RNA sequencing tracks highlighting the 3′-UTR profile differences for lengthened genes (A) or shortened genes (B) after 6 hours of PMA treatment of HL-60 cells overexpressing CFIm25 with respect to control for three biological replicates. The colors of the tracks represent OE-control (red) and OE-CFIm25 (blue). Proximal (P) and distal (D) poly(A) sites are indicated with arrows and chromosome co-ordinates and distance between the differentially used poly(A) sites are indicated at the top of each plot. The track labeled Annotated Poly(A) Sites gives the positions of known poly(A) sites from the polyA_DB database (Version V4.1s) identified in a variety of human cell types. For each of the browser plots, the y-axis value is reads per million (RPM) and the right side displays quantitative bar graphs reflecting the change in proximal poly(A) site usage with respect to the total read counts. An unpaired t-test was performed to determine the significance between the treatment groups and P value <0.05 was considered significant.

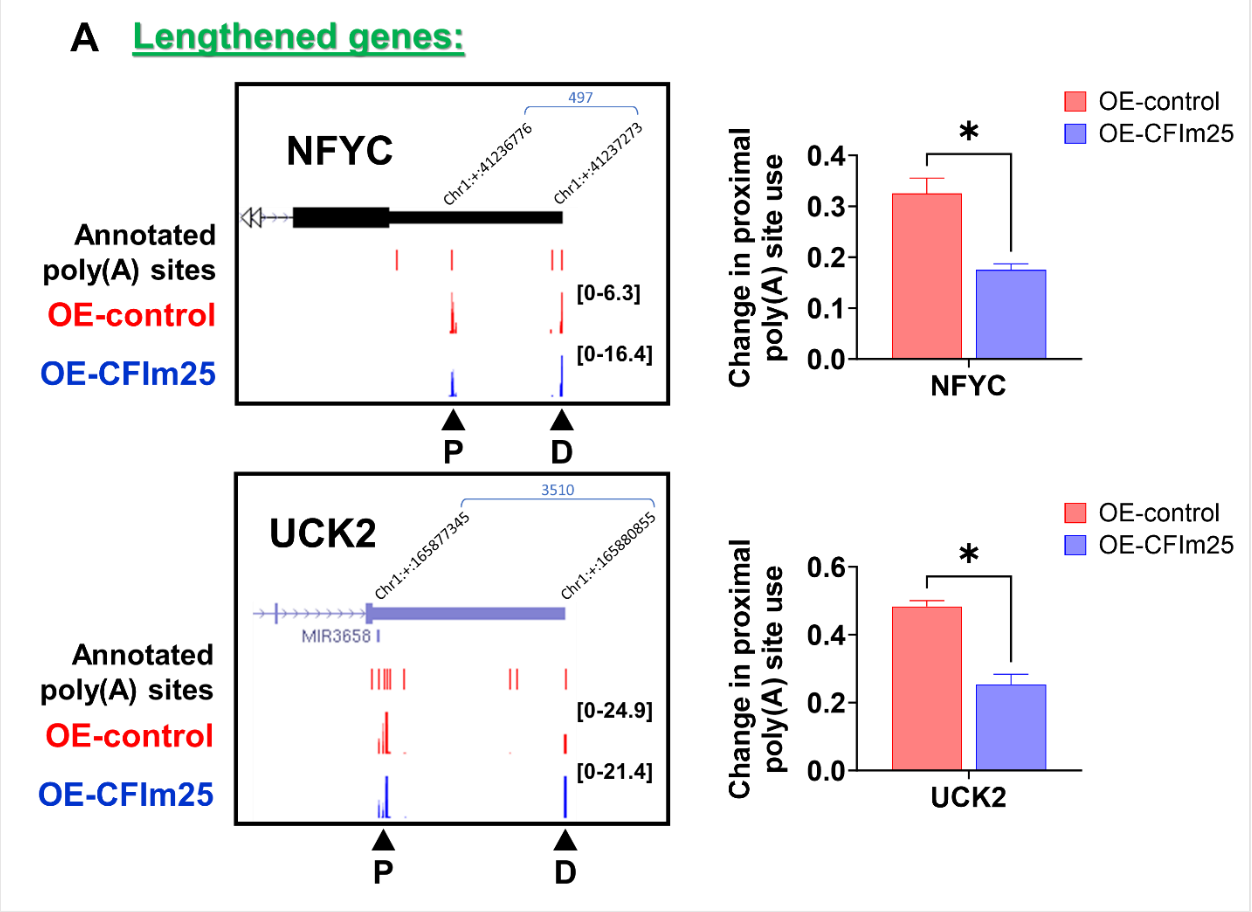

**B** Shortened genes:

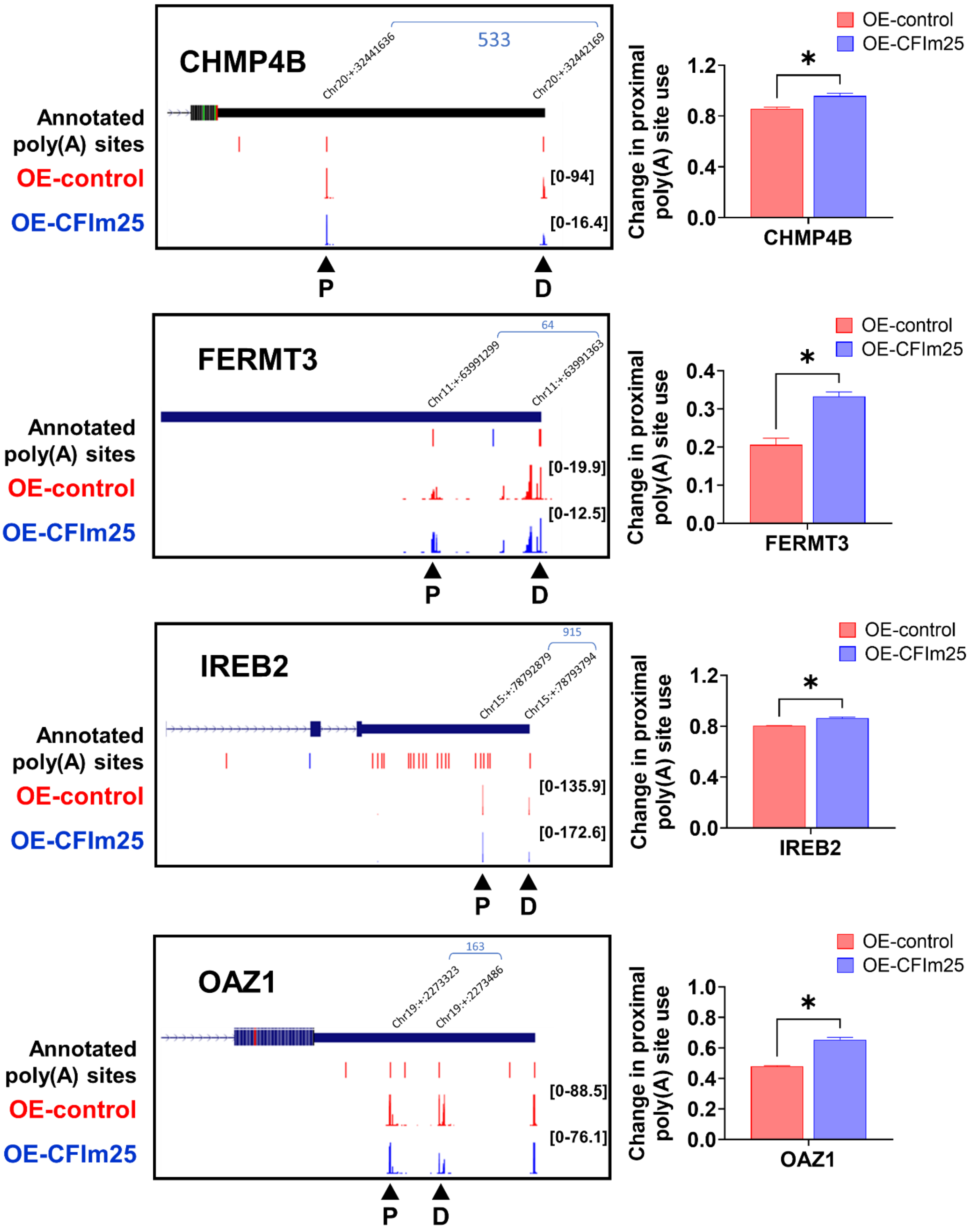

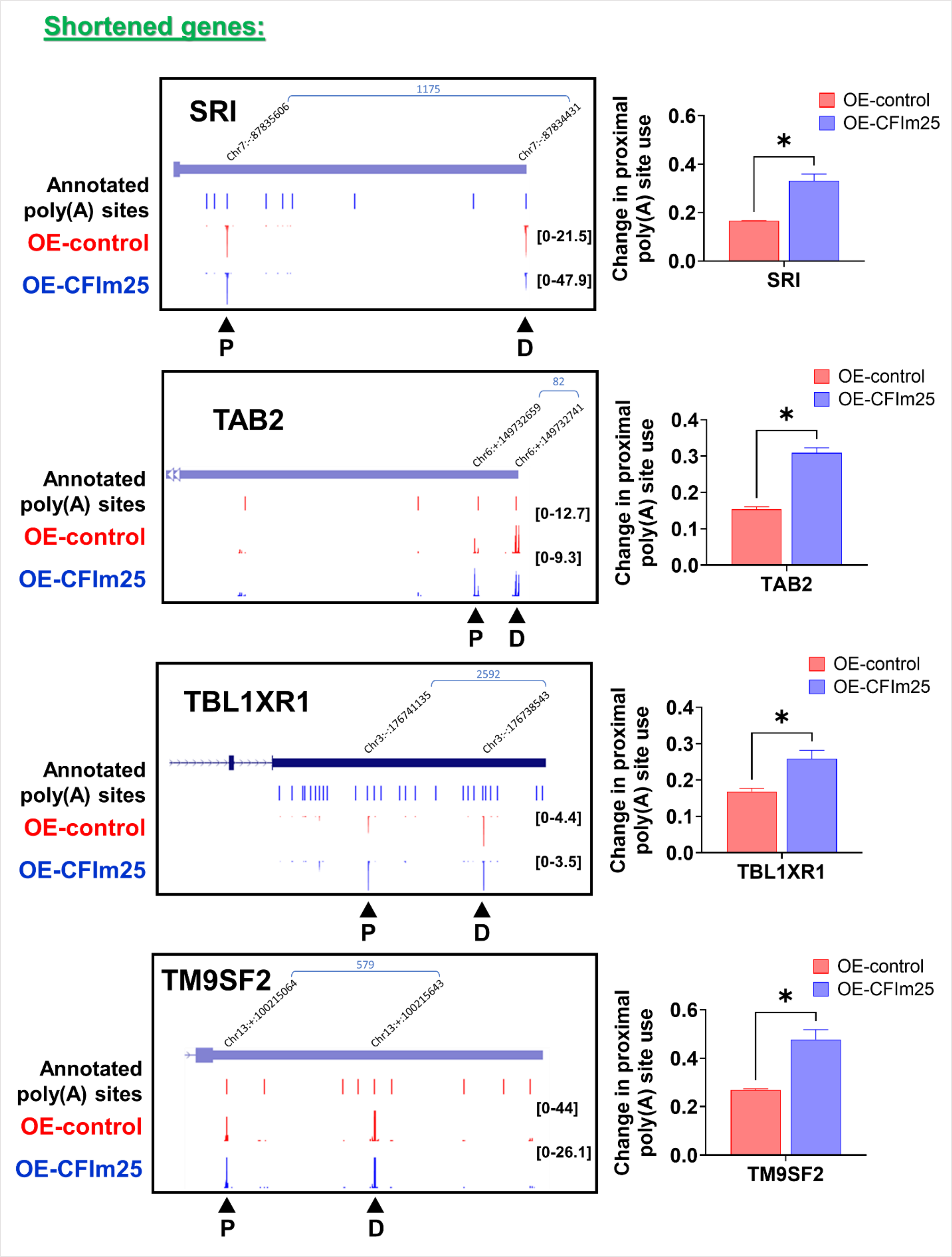


**Supplementary Fig 5. CFIm25 overexpression results in APA changes of specific genes and**

**altered protein expression in PMA-treated THP-1 cells. (A) APA and expression analysis of targets.** Real-time quantitative PCR (RT-qPCR)–based analysis of the expression and distal poly(A) site usage of different genes in THP-1 cells revealed by 3'Quant-seq as lengthened genes (*NFYC*, *UCK2)* or shortened genes (*CHMP4B*, *FERMT3*, *IREB2*, *OAZ1*, *SRI*, *TAB2*, *TBL1XR1* and *TM9SF2)*. The analysis was performed as described in Fig. 4. P value <0.05 was considered significant, where * = $P \leq 0.05$; **= $P \leq 0.01$; *** = $P \leq 0.001$. **(B) Protein levels of genes with APA changes.** Western blot analysis of proteins encoded by the shortened and lengthened genes as uncovered by 3'Quant-seq in THP-1 cells overexpressing CFIm25 with respect to control. GAPDH serves as the loading control.

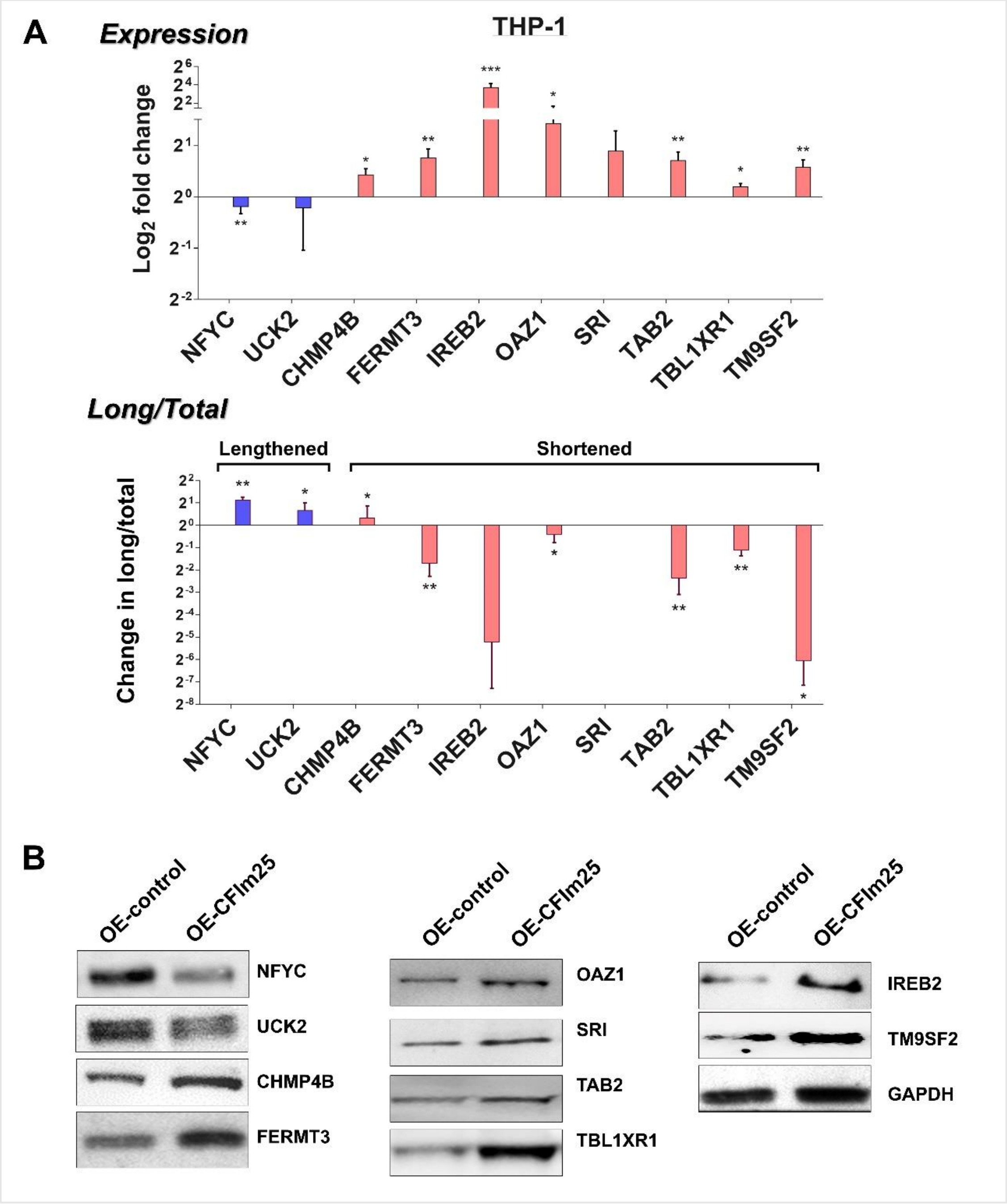

**Supplementary Fig 6. NF-κβ pathway inhibitor affects CFIm25 overexpression mediated differentiation. (A) CD38 levels in inhibitor treated HL-60 and THP-1 cells by flow cytometry.** Flow cytometry staining of surface marker CD38 at different time points during differentiation of control cells and cells treated with NF-κB inhibitor, where percentage represents cells positive for CD38. **(B) CD11b levels in inhibitor treated HL-60 and THP-1 cells by flow cytometry.** Flow cytometry staining of surface marker CD11b at different time points during differentiation of control cells and cells treated with NF-κB inhibitor, where percentage represents cells positive for CD11b. Gating for both is performed according to Supplementary Fig. 1C & D. Data is representative of at least three biological replicates.

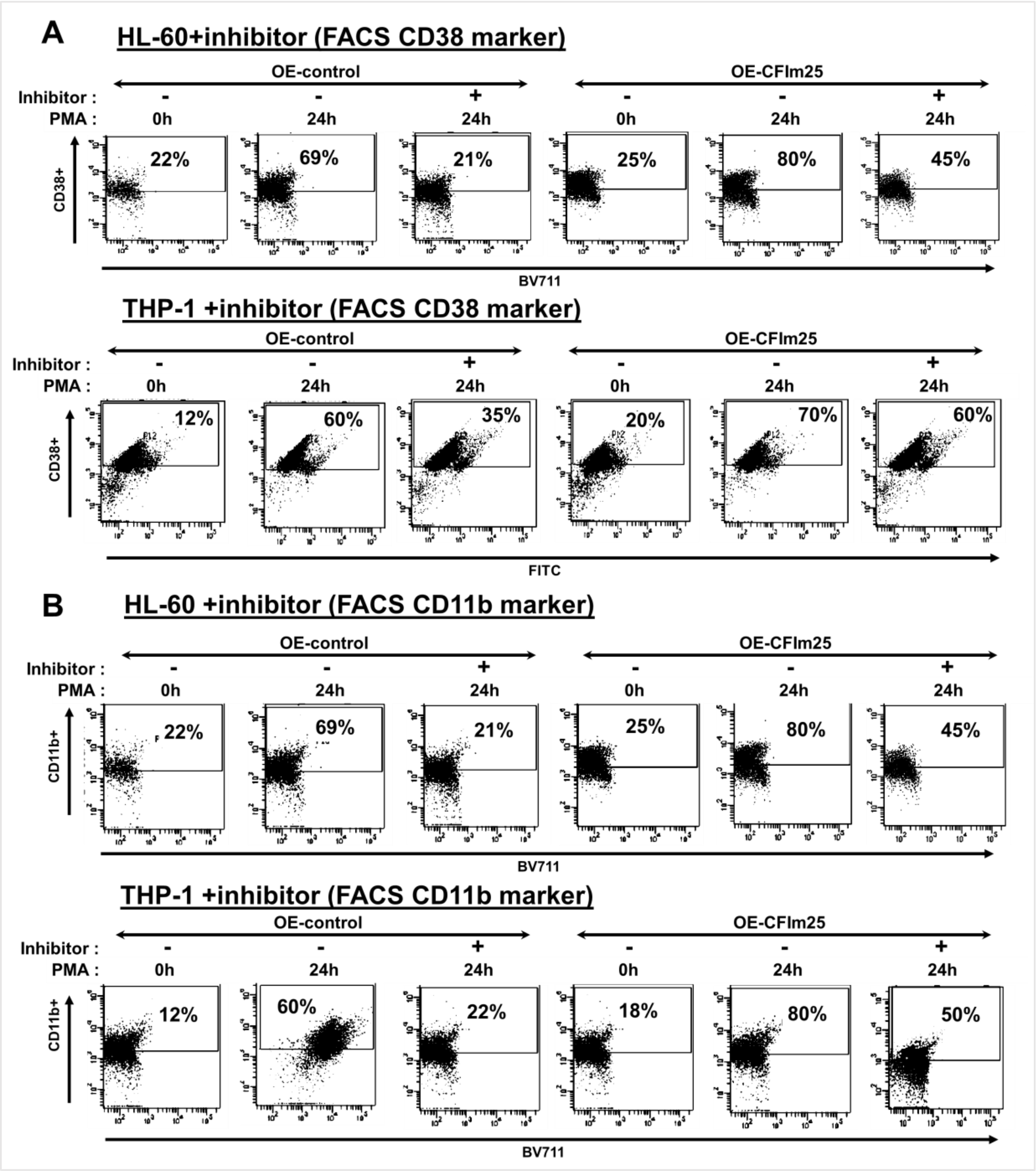
